# Conserved Laminar-Cellular Architecture Predicts the Spectrolaminar Motif

**DOI:** 10.1101/2024.03.27.587084

**Authors:** Aidan G. Mulvey, Maxwell J. Lichtenfeld, Hamed Nejat, Yihan (Sophy) Xiong, Brock M. Carlson, Blake A. Mitchell, Diego Mendoza-Halliday, Jacob A. Westerberg, Alexander Maier, Alvaro Duque, André M. Bastos

**Affiliations:** Department of Psychology, Vanderbilt University, Nashville, TN, USA; Department of Psychology, Yale University, New Haven, CT, USA; George Washington University School of Medicine and Health Sciences, Washington, District of Columbia, USA; College of Liberal Arts, University of Minnesota, Minneapolis, MN, USA; Salk Institute for Biological Studies, La Jolla, CA, USA; Department of Neuroscience, University of Pittsburgh, Pittsburgh, PA, USA; Department of Vision and Cognition, Netherlands Institute for Neuroscience, Royal Netherlands Academy of Arts and Sciences, Amsterdam, The Netherlands; MacBrain Resource Center (MBRC) - Department of Neuroscience, Yale University School of Medicine, New Haven, CT, USA; Vanderbilt Brain Institute, Vanderbilt University, Nashville, TN, USA

**Author notes:** **Corresponding Author:** André M. Bastos, Ph.D. 301 Wilson Hall, Vanderbilt University, Nashville, TN.

## Abstract

We recently identified the spectrolaminar motif, a layer-resolved pattern of local field potential (LFP) power, which is preserved across many areas of the primate cortex. It remains unclear which specific cytoarchitectonic features can explain the spectrolaminar motif. To investigate this, we performed histological and neurophysiological analyses of 11 brain areas spanning the macaque cortical hierarchy to examine the relationship between the laminar patterns of LFP power and the laminar distributions of cellular density for parvalbumin, calbindin, and calretinin interneurons, and pyramidal neurons. We tested whether cellular distributions are conserved and if they predict the corresponding variation in LFP power comprising the spectrolaminar motif. We found that the cytoarchitectonic and spectrolaminar data were intrinsically low-dimensional, with two-component patterns conserved across areas, and that the spectrolaminar motif could be predicted from these laminar cellular gradients. Critically, the regression coefficients mapping structure (cytoarchitecture) to function (spectrolaminar motif) were themselves highly conserved across areas, suggesting that a common cellular-to-spectral rule operates throughout cortex rather than a distinct rule per region. To evaluate this structure-function model, we tested it against post-mortem data from schizophrenia patients. We tested whether empirically-observed cytoarchitectonic changes from postmortem studies on inhibitory interneuron dysfunction in schizophrenia could be used to predict spectrolaminar alterations using our structure-function model. We found that they did, and that the functional changes match what is known from non-invasive neurophysiology in schizophrenia. This work moves beyond simple structure-function correlation to establish a predictive mathematical framework that informs healthy and aberrant brain circuitry.

## Introduction

We recently established neurophysiological evidence for a preserved spectrolaminar motif across 14 cortical regions in the primate neocortex.^1^ This work identified a conserved pattern of local field potential (LFP) power described by high-frequency gamma (30-150 Hz) activity peaking in cortical layers 2/3 and lower-frequency alpha/beta (8-30 Hz) activity peaking in deep layers 5/6. We also showed that layer 4 serves as a transition zone between alpha/beta and gamma power. These observations have been replicated in multiple independent labs, species, and additional cortical areas.^2–11^ In addition, this spectrolaminar motif is dependent on the orientation of cortex (i.e., the spectrolaminar pattern is flipped when recordings are superficial-to-deep versus deep-to-superficial), suggesting that the pattern is locally generated and not an artifact of distant volume conduction.^1,4,12^ However, it remains to be determined whether the spectrolaminar motif is supported by a preserved, underlying neuronal microcircuit. To address this, we collected histological data across many different cortical areas as well as laminar neural recordings from the same areas, to determine patterns of similarities or differences in the structure-function relationship.

Specifically, GABAergic interneurons are plausible candidates for drivers of the spectrolaminar motif. They are integral to the local circuit and the generation of theta, alpha/beta, and gamma frequency activity which comprise the spectrolaminar motif.^13–16^ In the primate neocortex, three distinct cell-types account for at least 90% of all interneurons with minimal colocalization and can be distinguished by their expression of calcium-binding proteins: Parvalbumin (PV), Calbindin (CB), and Calretinin (CR) positive interneurons.^15,17–22^ PV interneurons act proximally on pyramidal neurons located in layers 3/4 to provide the precise, rhythmic inhibition necessary to generate gamma oscillations.^14,23,24^ Superficial CB interneurons, which often colocalize with somatostatin (SST) interneurons in primates,^25,26^ can interact distally with pyramidal dendrites to modulate the beta frequency range.^14,27^ Superficial CR interneurons, which often co-express vasoactive intestinal peptide (VIP),^17^ have been shown to inhibit CB/SST interneurons and thereby disinhibit pyramidal cells.^28–30^ Also integral to the local circuit are glutamatergic pyramidal neurons, which can be immunohistochemically identified with the protein neurogranin (NG).^31,32^ Yet, while the laminar distributions and hierarchical patterns of interneuron subtypes have been mapped for some areas,^22,33,34^ and cell-type relationships with oscillatory activity have been established through optogenetics,^14,24,35^ these two bodies of work remain largely disconnected, especially for studies of the primate brain. Therefore, in this investigation we jointly studied multiple cell-types, cortical layers, and LFP spectral power to establish the linking principles between them.

An additional motivation of this work is to create structure-function models that may explain the functional consequences of cytoarchitectonic alterations observed in brain disorders. Specifically, aberrant GABA neurotransmission is hypothesized to cause neurophysiological disruptions and ultimately symptoms in disorders like schizophrenia.^36–38^ Most notably, PV interneuron deficits in cortical layers 3 and 4 have been robustly established through a recent quantitative meta-analysis of post-mortem data in human schizophrenia.^38^ Given the association between PV interneurons and gamma activity,^14,24^ deficits in PV are implicated in the selective weakening of gamma frequency activity that is characteristic of schizophrenia in task-evoked states.^39–41^ We hypothesize that *in-silico* modeling of these PV deficits will result in changes in gamma power reflective of that in schizophrenia.

To date, the link between neuronal architecture and laminar activity has been treated as a qualitative observation rather than a predictive principle.^42^ We address this gap by extending models of simple correlation. We instead utilized a multivariate mathematical framework to determine if the spectrolaminar motif is a predictable, emergent property of conserved cellular gradients. We aimed to uncover the cytoarchitectonic substrate for the spectrolaminar motif, thereby testing whether the structure-function relationships are conserved or heterogeneous across the cortical plane. We characterized the laminar distributions of PV, CB, and CR interneurons, and pyramidal neurons in 11 unique cortical areas, spanning the visual cortical hierarchy. We obtained matching laminar LFP data from these areas and quantified their area-specific spectrolaminar motifs. We employed principal component analysis (PCA) to characterize the prominent cytoarchitectonic and spectrolaminar gradients that exist in the primate neocortex. We tested whether these gradients were conserved across areas or unique to each. We then used partial least squares regression (PLSR) to create an accurate cytoarchitectonic model capable of predicting the spectrolaminar output of a given brain area. We hypothesized that the interaction between these inhibitory interneurons and excitatory pyramidal neurons within the cortical column would drive the observed spectrolaminar motif and that these circuit components were highly conserved between cortical areas. Establishing this structure-function relationship is critical, as many psychiatric disorders, like schizophrenia, have characteristic frequency aberrations^41^ and marked alterations in cellular composition.^38^ We therefore tested the structure-function model using the alterations in cellular composition observed in post-mortem schizophrenia data, to determine whether they resulted in frequency-specific changes matching previously known patterns in patient data obtained non-invasively.

## Methods

### Animal Procedures

All procedures were approved by the Vanderbilt University Institutional Animal Care and Use Committee in compliance with the regulations set by the Association for the Assessment and Accreditation of Laboratory Animal Care and followed the United States National Institutes of Health guidelines. Seven macaque monkeys were used in this study (**Table S1**). Two adult male bonnet macaques (*Macaca radiata*) were used: monkey Ca (19 years old and 7.4 kg) and monkey Bo (14 years old and 7.5 kg). Four adult male rhesus macaques (*Macaca mulatta*) were used: monkey Jo (11 years old and 12.0 kg), monkey Le (10 years old and 12.2 kg), monkey Sh (9 years old and 13.7 kg), and monkey St (10 years old and 12.1 kg). Monkeys Ca, Bo, Jo and Le were used for neurophysiology, and monkeys Bo, Le, Sh, and St were used for histology. We also used monkey B68 (*Macaca mulatta*; 18 years old and 8.18 kg) from the MacBrain Resource Center (MBRC; https://medicine.yale.edu/neuroscience/macbrain/) collection 6 for supplemental histology (**Table S1, S2**). Procedures regarding animal perfusion, immunohistochemical staining, and image capture in the MBRC are provided in detail elsewhere.^43^ Here, we provide such details for histology processed at Vanderbilt (monkeys Bo, Le, Sh, and St). See **Table S2** for the animal(s) used for each area.

### Neurophysiology: Data Collection

Monkey Ca was implanted with two MRI-compliant recording chambers: one over frontal areas (arcuate sulcus and dorsal/posterior end of principal sulcus) and another over the dorsal end of the superior temporal sulcus. Monkey Bo was implanted with one MRI-compliant recording chamber over the occipital cortex. Monkey Jo was implanted with one MRI-compliant recording chamber over the dorsal striate cortex. Monkey Le was implanted with one MRI-compliant recording chamber over the premotor cortex.

Monkeys were situated 57 cm from a PROPixx Pro projector (VPixx Technologies, Quebec, CA) at a resolution of 1920×1080 and a refresh rate of 120 Hz. Neurophysiological data were collected during the presentation of a full-field flash stimulus. Neural recordings were performed with either ‘Deep Array’ probes (Diagnostic Biochip, Inc., Glen Burnie, MD) or Plexon V-probes (Plexon Inc., Dallas, TX). Deep Array probes (monkeys Ca, Jo, Le) had 128 recording sites on each probe, with either 25 or 40 µm inter-contact spacing. Plexon V-probes (monkeys Bo, Le) had 32 recording sites on each probe organized as a linear array with an inter-contact spacing of 100 µm.

Acute penetrations were approximately perpendicular to the cortical surface and covered all cortical layers whenever possible. In this work, we assume that the spectrolaminar motifs from our recordings can be aligned to the histological data using a perpendicular mapping. This is well-supported by our previous work,^1^ in which we showed that the spectrolaminar motif is similar in probes with a perpendicular penetration as well as at larger angles of probe penetration relative to cortex (see Supplemental Methods). In this work, we also disregard differences between gyri and sulci, as our neurophysiology recordings and histological sampling include data from both. Recording areas were guided by MRI images and confirmed functionally with receptive field mapping on each recording day. Recordings were referenced to the guide tube in contact with both the probe and the dural tissue. All data were recorded with 30 kHz sampling using the Intan RHD recording system (Intan Technologies, Los Angeles, CA, US), then downsampled to 1000 Hz in offline preprocessing in MATLAB (The MathWorks, Inc., Natick, MA, US). LFP signal was extracted with a 4^th^-order Butterworth bandpass filter between 0.1 and 500 Hz.

Neurophysiological data collection was performed in primary visual cortex (V1), visual area three (V3), visual area four (V4), middle temporal cortex (MT), middle superior temporal cortex (MST), posterior inferotemporal cortex (TEO), dorsal premotor cortex (PMD), frontal eye field (8A), lateral prefrontal cortex (LPFC). Additional data was supplemented from publicly-available deposits from Mendoza-Halliday *et al.* (2024)^1^ in areas MT, MST, lateral intraparietal cortex (LIP), and LPFC (**Table S2**). This data was collected in monkeys Sh and St. Those recording procedures are described in detail elsewhere.^1^ All the areas used in the current study are diagrammed in Figure 1A and further detail is provided in Tables S1, S2.

**Figure 1.**
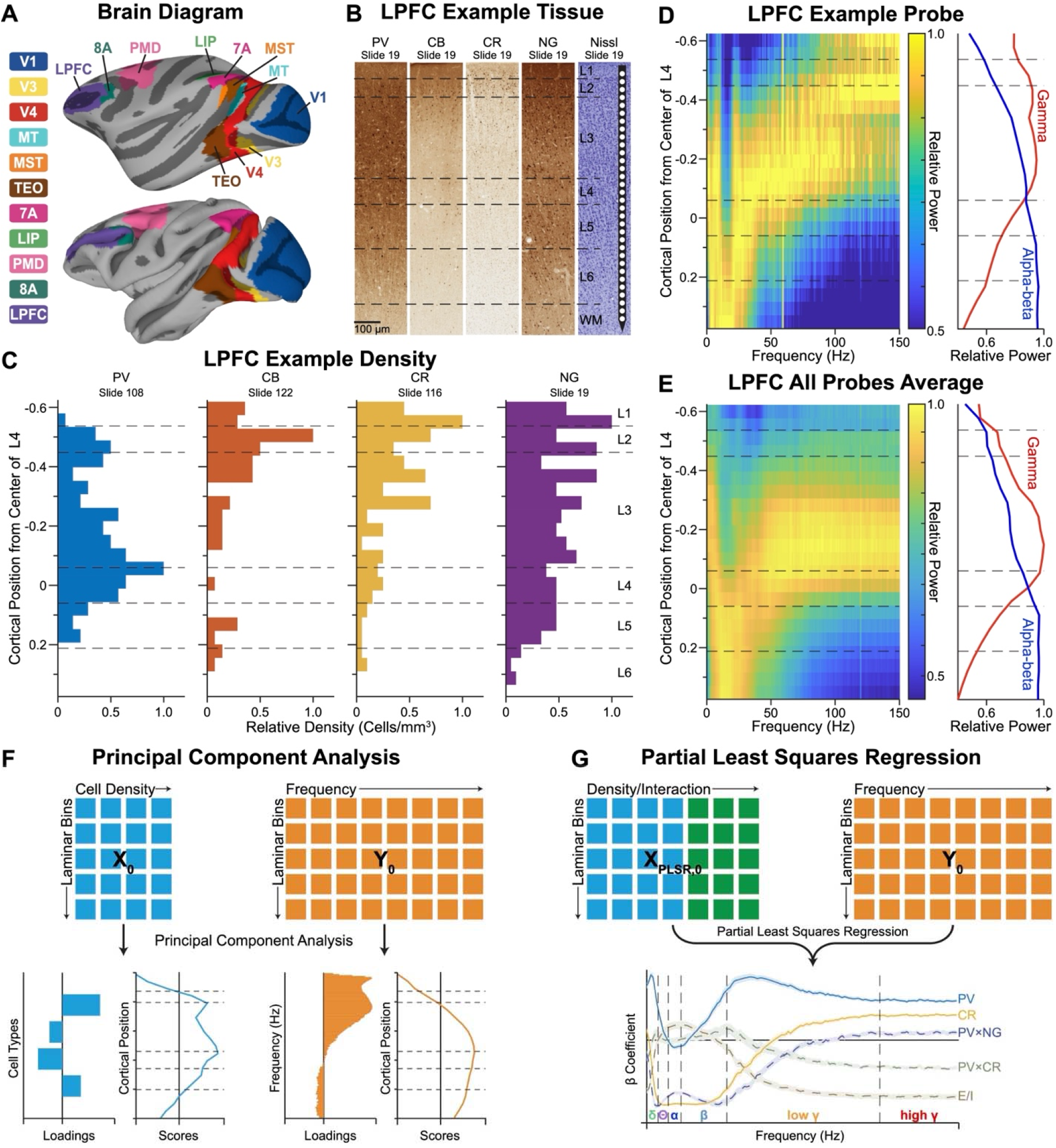
Histological and neurophysiological pipeline, and analytical framework. (A) Diagram of macaque brain with all areas included in the analyses, using the D99 Macaque Brain atlas in AFNI/SUMA (see Supplementary Methods). The inflated cortical surface (top) is shown to include layers visually blocked by gyri, as well as the semi-folded mid-thickness surface (bottom). Both are shown in the lateral, left hemispheric view. (B) Example “slabs” of LPFC from MacBrain Resource Center subject B68, slide 19. Layer boundaries are shown in dashed gray line, and a schematized 32-channel probe with contacts (white circles) is also shown to depict laminar recordings. A scale bar of 100 µm is shown as black line. (C) Example density distributions from single slabs from LPFC. Y-axes represent cortical depth relative to the middle of layer 4. Density shown in relative density (cells/µm^3^). (D) Single probe example of LPFC spectrolaminar graph. Y-axis represents cortical depth relative to the middle of layer 4. The x-axis represents frequencies from 1-150 Hz. The z-axis (color bar) represents relative power. Average alpha-beta power (8-30 Hz) shown in blue line and average gamma power (31-150 Hz) shown in red line. (E) LPFC spectrolaminar graph averaged across all probes (n = 38). (F) Schematized principal component analysis framework with the cytoarchitectonic (**□**_0_) and spectrolaminar (**□**_0_) datasets. Output of each PCA includes loadings and scores, where loadings indicate the strength of the contribution of each cell-type or frequency, and the scores indicate the laminar localization of the component. (G) Schematized partial least squares regression framework. Cytoarchitectonic dataset (**□**_PLSR,0_) includes the cell types and cell type combinations. Output includes regression coefficients (β) of each cell type and cell type combination as a function of frequency.

### Neurophysiology: Data Analysis

All statistical analyses were conducted in MATLAB (version 2024a, The MathWorks, Inc., Natick, MA, US). Power analysis was performed on a one second, peri-stimulus time window (500 ms pre-stimulus to 500 ms post-stimulus). We used the *freqanalysis* function with *mtmfft* parameter from the open-source FieldTrip toolbox.^44^ This implements a multitaper spectral estimate 68. We used 2 Hz smoothing in the spectral domain. Power was calculated on individual trials and then averaged across trials. We then obtained the relative power maps for each probe separately (equation 1), where *c* is each channel on the probe and *f* is each frequency (1–150 Hz). For each probe this resulted in a 2-dimensional matrix, with frequency on the x-axis and channels on the y-axis (**Figure 1D**). Thus, at each frequency, every channel had an intensity between 0 and 1, where values of 1 indicate the channel that had the highest power at that frequency. We interpolated over 60 Hz to remove noise using *interp1*. Finally, we averaged across probes to obtain a single spectrolaminar motif per area (**Figure 1E**).

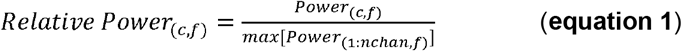

### Histology: Perfusion Surgery

At Vanderbilt, monkeys were anesthetized with ketamine (7.5 mg/kg IM) and dexmedetomidine (0.015 mg/kg IM). Lethal sodium pentobarbital solution (40 mg/kg IV or greater) was started immediately after the placement of probe penetration angle reference noodles. Monkeys were perfused transcardially with 30% phosphate buffered saline, followed by 4% paraformaldehyde, and finally 4% paraformaldehyde with 10% sucrose. The whole brain was stored in sucrose phosphate buffered saline until ready for sectioning and slicing. Perfusion surgery for monkeys Sh and St was performed at Massachusetts Institute of Technology (MIT) and is described in detail elsewhere.^1^ However, histology for monkeys Sh and St was subsequently processed at Vanderbilt University and, thus, is described here. The brain was sectioned along the hemispheric midline and then cut into separate blocks for each chamber. These MRI-guided cuts were in plane with the probe penetration angles (reference noodles), and so they were not sliced in a canonical direction. Blocks were sliced at 40 or 50 µm on a freezing microtome. Slicing thickness was uniform within each animal, and thickness was selected based on the stability of the tissue when being sliced. Slicing thickness was recorded and is included in calculations of density.

### Histology: Immunohistochemistry

All odd-numbered sections were Nissl stained. Sections to be processed for markers of PV, CR, and CB were each selected as one in a series of twelve, resulting in twelve sections between consecutive stains of the same type (e.g., PV to PV section). Sections were processed for PV, CR, and CB using the protocol described by Ichinohe *et al.* (2003).^45^ We used mouse monoclonal anti-PV (RRID:AB_10000343), mouse monoclonal anti-CB (D28k; RRID:AB_10000347), and mouse monoclonal anti-CR antibodies (RRID:AB_10000320). All antibodies used were produced by Swant, Bellinzona, Switzerland.

Between each independent step, the sections were rinsed three times in a wash solution composed of 0.1 M phosphate-buffered saline (PBS; pH 7.2) with 0.5% Triton X-100. Sections were incubated for one hour at room temperature in a blocker solution composed of 0.1 M PBS (pH 7.2) with 0.5% Triton X-100 and 5% normal horse serum. The sections were then incubated for 48 hours at 4°C in their respective primary antibodies within a solution of blocker. In the following step, sections were incubated in a blocker solution containing biotinylated horse anti-mouse IgG (1:200; RRID:AB_2687893; Vector, Burlingame, CA, US) for 90 minutes at room temperature. Next, sections were incubated for 90 minutes at room temperature in Avidin-Biotin Complex (ABC) solution (one drop each of reagent A and B per 7 mL of 0.1 M PB, pH 7.2; ABC kits, Vector Labs, Burlingame, CA, US). Immunoreactivity was visualized through development of the section in diaminobenzidine histochemistry with 0.03% nickel ammonium sulfate. Sections were mounted and secured to glass slides for imaging.

### Histology: Image Capture and Processing

We imaged each Nissl-, PV-, CB-, and CR-stained slide using a Zeiss Axio Imager.M2 microscope (Zeiss, Jena, Germany). The Nissl-stained slides were taken at a magnification of 1.25x, which corresponds to a pixel to space conversion factor of 8 µm per pixel. PV-, CB-, and CR-stained slides were taken at a magnification of 10x, which corresponds to 1.03 µm per pixel. Images of NG-stained tissue and supplemental tissue stained for inhibitory cell markers (PV, CR, and CB) for areas that were not collected at Vanderbilt were obtained from the MBRC (**Table S2**). MBRC slides were imaged at 20x and subsequently scaled down to 10x for ease of processing and congruence with tissue processed and imaged at Vanderbilt.

Once imaged, brain areas were located by cross-referencing anatomical landmarks found in the images with both MRI data from the particular animal and *A Combined MRI and Histology Atlas of the Rhesus Monkey Brain in Stereotaxic Coordinates*.^46^ We collected histological data from areas V1, V3, V4, MT, MST, LIP, TEO, PMD, 7A, 8A, LPFC (**Figure 1A, Table S2**). After isolating areas, we sampled at least three unique slides for each cell type. Tissue was then oriented to align the cortex to a perpendicular position relative to a line from the top of cortex (pia, top of layer 1) through the bottom of cortex (white matter, bottom of layer 6). We created rectangularly cut samples spanning a uniform 200 µm in width and a length measuring the entirety of the laminar column (n = 15 “slabs” per area per stain type; see **Figure 1B** for example slabs in LPFC).

### Histology: Cell Counting and Measurements

Cell counting was performed using the Fiji ImageJ software.^47^ Cells were counted using a semi-automated technique with intensive human review by two independent reviewers (M.J.L. and A.G.M.). Slabs were loaded into ImageJ and contrast and brightness were adjusted to clearly distinguish the darkly stained cell bodies from the lighter background. The human reviewer reviewed each contrasted slab to eliminate noise (e.g., darkly stained non-cell bodies) and increase signal (e.g., isolate more lightly stained cell types that were excluded by the contrasting steps). We then applied the built-in ImageJ particle detecting tool to detect the cell bodies, and automatic measurements were taken from the top of the sample and from the left edge of the sample to the center of the cell bodies in the unit of pixels.

### Histology: Laminar Distribution Calculation

Laminar boundaries and height of cortical columns were identified in adjacent Nissl slabs. To standardize measurements within an area, all measurements were converted into “standardized space” defined by the center of layer 4 as zero. Locations more superficial than layer 4 (supragranular layers) were negative, and locations deeper than layer 4 (infragranular layers) were positive. To convert cell location measurements to this relative space, we first extracted the location of the center of layer 4 from the top of the cortex using Nissl slabs. We then subtracted this value from the cell depth measurement and divided it by the total cortical height of the Nissl slab. The resulting measurements are in units of proportion of cortical depth away from the center of layer 4, with the total length from the top of cortex to the bottom being one. For example, if a cell was measured at a depth of 250 µm from the top of layer 1 with a total cortical depth of 2000 µm and the depth of the middle of layer 4 being 1000 µm the conversion would go as such:

1. 250 µm – 1000 µm = −750 µm (distance from the middle of layer 4 to that cell)
2. -750 µm / 2000 µm = −0.375 (the cell is 0.375 of the total cortical depth, superficial of the middle of layer 4)

To create laminar distributions of cell counts, we created 100 µm bins across the entire length of a cortical area (e.g., if cortex was 2 mm, we had 20 bins). We then created our laminar profiles using the *histcount* function, assigning the cortical position of a cell to a bin. We calculated the volume of each bin by multiplying the bin height (100 µm) by the slab width (200 µm) by the slice thickness (40/50 µm). We then divided the cell counts by the volume to obtain cellular density in units of cell/µm^3^, which was used throughout the subsequent analyses (**Figure 1C**). We repeated this 15 times (n = 15 “slabs”). To easily cross-compare MBRC tissue to other cell types and to spectrolaminar data, we established the mean laminar bounds for each area, averaging all the laminar boundaries for the 15 slabs within an area. We also correlated the density distributions across histology subjects to confirm that distributions are similar across animals (**Figure S2A, B**).

To align cytoarchitectonic and spectrolaminar data, we assumed that the laminar position of layer 4 could be identified using the cross-over point between alpha-beta and gamma power, as demonstrated with histological verification by Mendoza-Halliday *et al*. (2024)^1^ and Major *et al*. (2026).^2^ Cortical boundaries were defined using histological markers to measure the distance (in mm) from the middle of L4 to the top and bottom of the cortical column. Probe channel locations were then converted in absolute distance in millimeters, scaling the probe coordinates to match the anatomical boundaries relative to the middle of layer 4, which aligned cytoarchitectonic and spectrolaminar data.

### Histology: Uniform Space Transformation

We used a piece-wise linear rescaling function on the cytoarchitectonic and spectrolaminar datasets to convert both to a uniform space, allowing the comparison across areas. We calculated the mean proportions from each area to be set as our uniform boundaries. We determined the cortex to be divided 60% above the middle of layer 4 and 40% below by taking the average bounds across all areas. We therefore chose −0.6:0.4 with steps of 0.05, creating 21 cortical bins. From the native space, we interpolated the top half of cortex to [-0.65, 0] and the bottom half of cortex to [0 to 0.45) independently, generating 23 bins. We excluded the first and last bins to avoid any surface level artifacts from interpolation, creating 21 total cortical bins. Thus, all areas were converted into this standardized, 21 cortical bin space that had proportional laminar boundaries from −0.6 to 0.4.

### Statistical Analyses: Preparation of Datasets

Here, we refer to the cytoarchitectonic (histology) dataset as **□** and the spectrolaminar (LFP) dataset as **□**. **□** is a 21×4×11 matrix, where 21 is cortical bins, 4 is cell types, and 11 is number of areas. **□** is a 21×150×11 matrix, where 21 is cortical bins, 150 is each frequency, and 11 is number of areas.

To assess whether the laminar organization of the cytoarchitecture and spectrolaminar motif is preserved across areas, we computed Pearson correlations between every pair of areas (11 areas, 55 pairs) and summarized conservation as the mean pairwise *r*. For each cell type, correlations were computed between laminar distributions across areas. For the spectrolaminar motif, each area’s 21×150 power spectrum unrolled into a single 3150×1 vector. Pearson correlations were then computed between these vectors across areas. The same analysis was repeated for each isolated frequency band (delta, theta, alpha, beta, low gamma and high gamma). Significance was assessed by permutation testing (N = 5,000 permutations). *P*-values were reported as the proportion of permutations with the mean pairwise *r* equivalent to or exceeding the observed value.

We then calculated the mean distribution of cell density across all areas (n = 11) for each cell type (**□**_0_), creating a **N**×**P** matrix, where **N** is 21 (cortical bins) and **P** is 4 (cell types). Similarly, we took the mean relative power at each cortical bin for each frequency across all included areas (**□**_0_), producing a **N**×**Q** matrix, where **N** is 21 and **Q** is 150 (frequencies). This allowed the generation of a “prototypical” cortex, described by the mean distribution of each cell type and mean relative power at each frequency bin. We applied a z-score to the feature dimension (cell-types or frequencies) of each dataset, which mean-centers the data and prepares it for further statistical analyses.

### Statistical Analyses: Principal Component Analysis

To understand the latent composition of the **□**_0_ (structural cytoarchitecture) and **□**_0_ (functional spectrolaminar motif) matrices, we performed principal component analysis (PCA; **Figure 1F**). Using the *pca* function on the **□**_0_ and **□**_0_ matrices separately, we obtained the principal component coefficients (variable contributions), principal component scores, eigenvalues, and percentage of explained variance. We retained the number of principal components (PCs) necessary to account for at least 90% of the cumulative variance. To determine the stability of the coefficients, we applied a bootstrapping procedure (N = 5,000 resamples).^48^ We calculated the bootstrap ratios (BSR) for each coefficient of **□**_0_ and **□**_0_ to determine which coefficients stably contribute to the observed PCs.^49^ We used a conservative bootstrap threshold of 2.5 (p ≈ 0.01), such that a |BSR| ≥ 2.5 is stable (p < 0.01).

While PCA was done on the global (averaged across areas) matrices **□**_0_ and **□**_0_, we also wanted to understand how well each individual area conformed to the identified global patterns. We calculated PCs 1 and 2 for each area (**□***_i_* and **□***_i_*, where *i* is each area), then performed Procrustes analysis,^50^ using the function *procrustes*, on the PC 1 and 2 scores for the global **□**_0_ and **□**_0_ laminar patterns against each individual area. This gave us the Procrustes distance *d*, which is a number between 0 and 1, with 0 being extremely close (i.e., similar) and 1 being extremely far (i.e., dissimilar) between the global patterns and each area’s pattern. We also permuted (N = 5,000 permutations) each area’s cytoarchitectonic and spectrolaminar data to obtain null distributions that establish a critical cutoff value for the Procrustes distance. We established a critical *d* cutoff by calculating the 1st percentile of the null distributions of permuted distances for **□**_0_ and **□**_0_, yielding d(**□**) = 0.756 and d(**□**) = 0.752. Thus, any area that exceeded these bounds significantly differed from the identified global laminar patterns.

### Statistical Analyses: Partial Least Squares Regression

Partial Least Squares Regression (PLSR) is a multivariate, data-driven regression technique widely used in neuroimaging to predict one high-dimensional dataset from another. PLSR is particularly useful when the number of observations (**N**) is limited, the number of dependent or independent variables (**P** and **Q**) exceeds the number of observations, or in the case of multicollinear data.^51–56^ PLSR extracts a small number of latent variables that maximize the covariance between the predictor (cytoarchitecture, **□**_0_) and the response (spectrolaminar motif, **□**_0_), while avoiding overfitting through cross-validation. Unlike ordinary least squares, PLSR is particularly useful in small-N, high-dimensional datasets like the present one. We applied it here to determine if the spectrolaminar output of cortex (**□**_0_) is a predictable property of conserved cytoarchitectonic distributions (**□**_0_).

To more completely understand the role of neuron types in predicting the laminar power spectrum, we added 6 cell type combinations via the iterative multiplication of each z-scored cell type distribution (i.e., PV×CB, PV×CR, PV×NG, CB×CR, CB×NG, CR×NG) and 1 proportion term, which is the z-score of the pyramidal cell distribution divided by the cumulative distributions of the inhibitory cell types (i.e., approximating E/I balance over layers). Thus, **□**_P_ is a 21×11×11 matrix, where 21 is cortical bins, 11 is cell types and their combinations, and 11 is number of areas. **□**_P0_ is the average of **□**_P_ across the area dimension (**Figure 1G**). **□**_0_ is the matrix of relative power averaged across all areas. We used the *plsregress* function on **□**_P0_ and **□**_0_. We then calculated the predicted spectrolaminar motif (**□**_PRED_) using the formula **□**_PRED_ = **□**_P0_×β, where β is the 11×150 matrix of standardized regression coefficients.

To determine the robustness of the structure-function model, we performed a Leave-One-Area-Out cross-validation (LOOCV), where we excluded one area from **□**_P_ and from **□**, such that each area is systematically excluded from the training set. In each round of exclusion, we computed the Pearson correlation (*r*_PRED_) between the observed (observed test set, **□**_TEST_) and predicted spectrolaminar motif (**□**) of the excluded area (predicted test set, **□**_PRED,OUT_) as the average of per-frequency correlations. To assess internal robustness and predictive strength of the model, we calculated Q^2^ across LOOCV iterations (equation 2), where <u>*Y*</u> is the mean of the training set **□**.^57^ Q^2^ is the cross-validated equivalent of R^2^, and it measures the fraction of the held-out area’s observed spectrolaminar motif that the prediction accounts for.

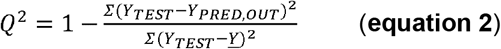

The standard error of β and *r*_PRED_ were calculated across LOOCV iterations. The variances associated with *r*_PRED_ and Q^2^ allowed us to describe the conservation of the predictive relationship and the predictive strength, respectively, across areas. Within each frequency band, *r*_PRED_ and Q^2^ were averaged across frequencies for each held-out area, and significance was assessed using a two-sided Wilcoxon signed-rank test against zero across areas. This was done to determine the significance of each frequency band. We evaluated the model’s predictive performance as a function of the number of components. In PLSR, components are the weighted combination of the 11 cytoarchitectonic predictors chosen to maximally covary with the spectrolaminar data. Latent components are created by the model to prevent overfitting across 11 correlated predictors by reducing redundancy and maintaining a shared signal. A 4-component model was selected as it had the strongest *r*_PRED_ and Q^2^ value without overfitting. These 4 components explained a combined 97.60% and 90.97% of the variance in the cytoarchitectonic and spectrolaminar datasets, respectively, suggesting these components accurately represent the datasets. Overall significance of the PLSR model was assessed using Fisher’s combined probability test,^58^ after permuting (N = 5,000 permutations) and recalculating Q^2^ on the permuted PLSR results.

We next collapsed the continuous 150 frequencies into six discrete frequency bands (delta [1-3 Hz], theta [4-7 Hz], alpha [8-12 Hz], beta [13-30 Hz], low frequency gamma [31-90 Hz], high frequency gamma [91-150 Hz]). We repeated the PLSR-LOOCV procedure on the six frequency bands. While this is less informative than the continuous spectrum, this provides a more concise interpretation of the structure-function relationship (i.e., one β per frequency band).

To determine area-wise robustness, we performed PLSR on each individual area, yielding a β_i_ matrix for each area. We then compared the predicted spectrolaminar motif (**□**_i_ = **□**_Pi_×β_i_) of one area with another area’s observed spectrolaminar motif (**□**_0_) using a 2-dimensional Pearson correlation (*corr2*). This was done to determine if one area’s cytoarchitecture could predict another area’s spectrolaminar motif or if each area was unique and thus unable to predict the spectrolaminar output. We also correlated β_i_ across areas to determine if the mathematical translation between cytoarchitecture and spectrolaminar motif (β_i_) is ubiquitous or distinct in each area. We assessed the significance of this analysis via a permutation test (N = 5,000 permutations), where we shuffle the laminar bins in cytoarchitecture to break the structure-function relationship.

### Statistical Analyses: Schizophrenia Perturbation Analysis

To explore clinical relevance with a causal test of the PLSR model, we conducted a targeted *in-silico* perturbation analysis. GABAergic interneuron deficits are a well-replicated pathological feature of schizophrenia. We used the Hedges’ *g* effect sizes for PV, CB, and CR interneuron density from a recent meta-analysis on GABAergic interneuron pathology in schizophrenia^38^ to selectively knock-down these interneurons by the reported effect size across cortical layers (see Supplementary Methods). We then re-computed the cell type combinations using the perturbed cytoarchitecture and calculated the predicted spectrolaminar output based on β from the PLSR-LOOCV model, which assumed that, even in an altered model, the relationship between cell types and spectral power remains constant. We calculated the percent change between the control and perturbed model spectrolaminar motifs. This was repeated for all 11 areas, and we used a Wilcoxon signed rank test to determine significance of the percent change at each frequency across areas.

## Results

### Cytoarchitectonic and Spectrolaminar Datasets

Each cell type exhibited conserved patterns across areas (**Figure 2A**). PV was marked by a bimodal peak in layers 3 (L3) and 4, with the L4 peak appearing more prominent. CB and CR had similar superficial unimodal peaks, though some early cortical areas (V1-MST) had a secondary peak in deep L3. NG displayed higher variability across areas, mostly characterized by a primary peak in superficial L3 and a secondary, attenuated peak in deep L5/6. Using a pair-wise Pearson correlation and permutation test, we determined that all cell type laminar distributions were conserved across areas (**Figure 2B**). CR interneurons were the most conserved (mean *r* = 0.81, *p* < 0.001), then CB (mean *r* = 0.70, *p* < 0.001), PV (mean *r* = 0.64, *p* < 0.001), and NG (mean *r* = 0.53, *p* < 0.001).

**Figure 2.**
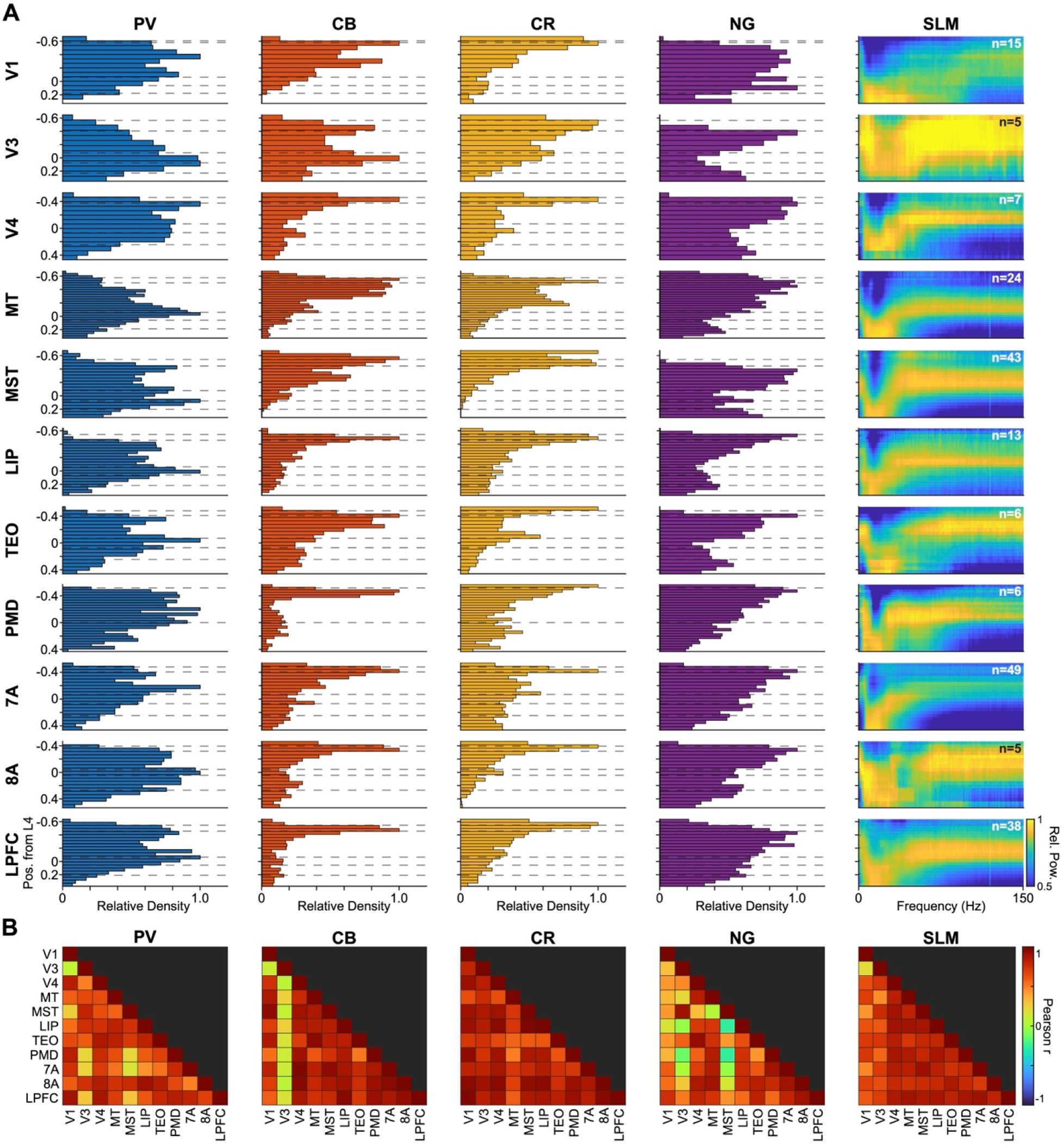
Laminar Profiles of Cytoarchitectonic and Spectrolaminar Motifs Across Areas. Relative density distributions of PV (blue), CB (orange), and CR (yellow) interneurons, and NG (purple) cells across areas. Spectrolaminar motif graphs are also shown across areas, with the number of recordings per area indicated in the top right of each spectrolaminar graph. Relative power of each graph ranged from 0.5 to 1. Laminar depth is given as the proportion of cortex from the middle of layer 4. (B) Pearson *r* values from correlations of density distributions and spectrolaminar motif across areas.

We then characterized the spectrolaminar patterns across the 11 brain regions (**Figure 2A**). Replicating Mendoza-Halliday *et al.*, we identified a highly conserved pattern in the spectrolaminar dataset, characterized by a laminar gradient of increasing alpha/beta power from superficial to deep layers, peaking in L5/6, and a laminar gradient of increasing gamma power from deep layers to middle layers, peaking in L3. Using a pair-wise Pearson correlation on the entire two-dimensional power spectrum and a permutation test, we determined that the spectrolaminar motif was significantly conserved across cortical areas (mean *r* = 0.76, *p* < 0.001; **Figure 2B, Figure S3**). High gamma power (91-150 Hz) was the most conserved (mean *r* = 0.89, *p* < 0.001), then beta (13-30 Hz; mean *r* = 0.78, *p* < 0.001), alpha (8-12 Hz; mean *r* = 0.63, *p* < 0.001), low gamma (31-90 Hz; mean *r* = 0.63, *p* < 0.001), delta (1-3 Hz; mean *r* = 0.52, *p* < 0.001), and theta (4-7 Hz; mean *r* = 0.49, *p* < 0.001). The average of the cellular density distributions and spectrolaminar motifs across areas are shown in **Figure 3**.

**Figure 3.**
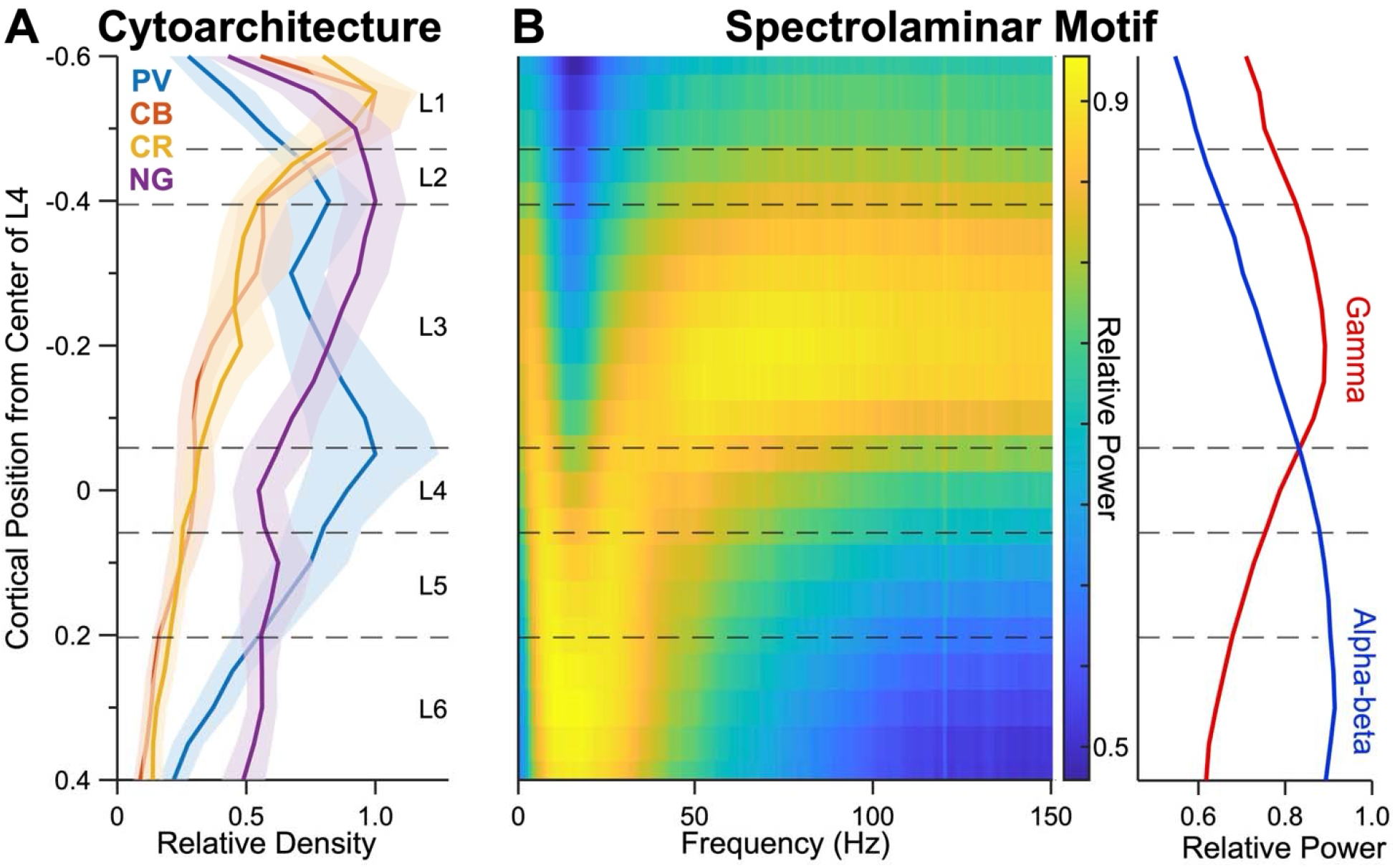
Prototypical Cytoarchitecture and Spectrolaminar Motif Across Areas. (A) Mean density of PV, CB, and CR interneurons and NG neurons across areas shown in solid lines ± 2 SEM. Variance is calculated across areas. (B) Mean spectrolaminar graph across areas and mean relative power in the alpha-beta (blue) and gamma (red) bands. Both shown as functions of cortical position from the middle of layer 4.

Previous work established that the spectrolaminar motif is robust across other primate species and independent research laboratories.^1,2^ To ensure that our results were reproducible, we compared the spectrolaminar motif across recording sessions, animals, and sites/institutions. We confirmed that the spectrolaminar motif is highly similar across all sessions, animals, sites, and brain regions tested (**Figure S1A, B**).

### PCA Yields Two Components in Both Structure and Function

We next wanted to understand whether the complex cytoarchitectonic and spectrolaminar motifs could be decomposed into simpler, orthogonal components. We applied principal component analysis to the average cellular distributions and spectrolaminar motif across areas (**Figure 3A, 3B**). We identified two principal components from both the cytoarchitectonic and spectrolaminar datasets, explaining a combined 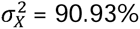 and 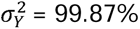 of the total laminar variance in cellular density and frequency-wise spectral power respectively (**Figure 4A-C**). PC1_X_ described the superficial CB/CR laminar gradient (σ^2^ = 62.47%; **Figure 4A**). CB and CR were the primary contributors to this PC, but NG was found to be stable after bootstrapping (all |BSR| > 2.5; **Figure S4**). PC1_X_ peaked in L1/2, then steadily declined into L4-6, where it troughed in deep L6. PC2_X_ described the L3/4 PV gradient (σ^2^ = 28.46%; **Figure 4A**). This PC peaked at the L3/4 border and only PV stably contributed to this axis (|BSR| > 2.5). Taken together, these PCs suggest that cytoarchitecture is governed by two orthogonal axes captured by PC1_X_ and PC2_X_.

**Figure 4.**
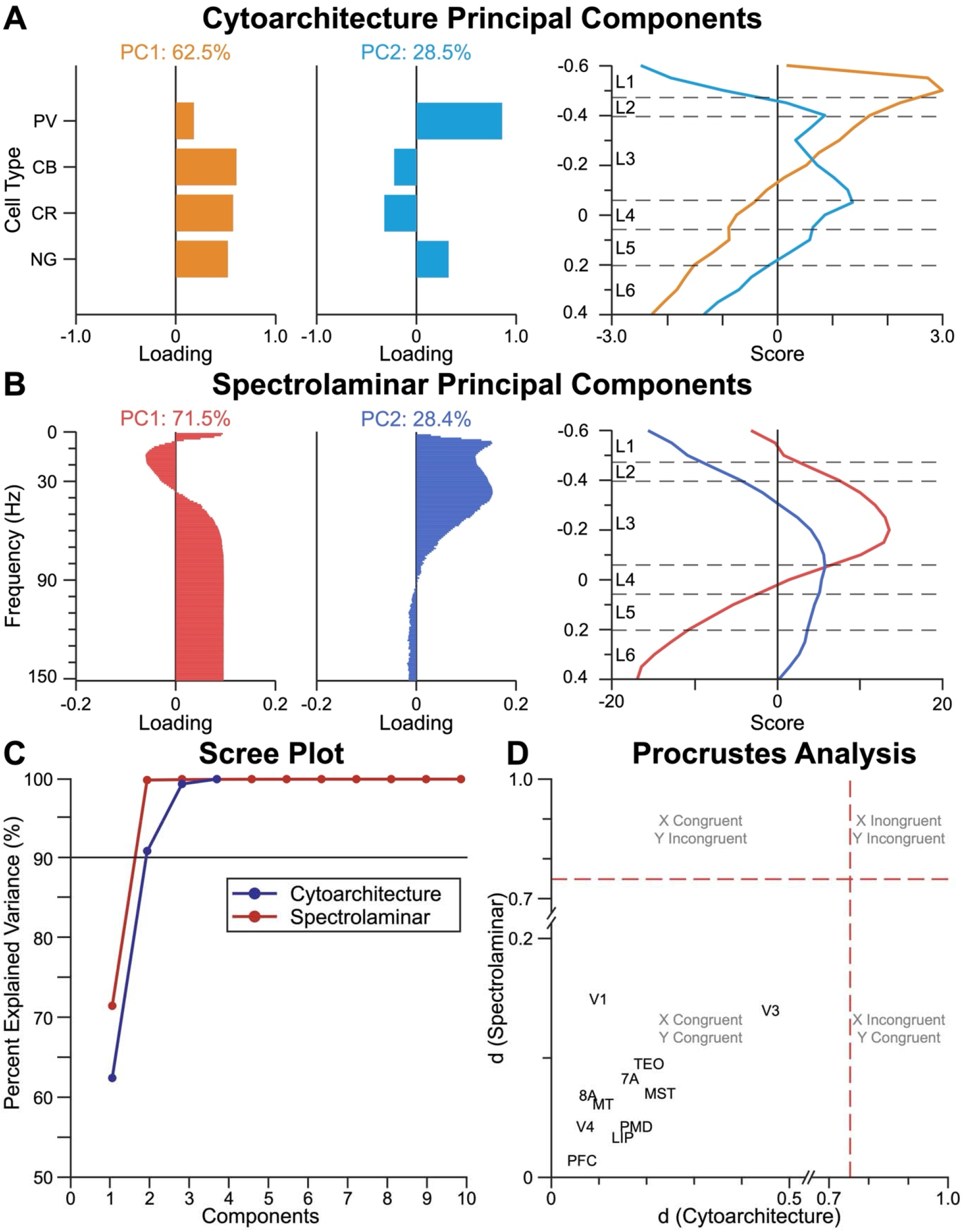
Principal Component Analysis of Cytoarchitectonic and Spectrolaminar Motifs. (A) Primary and secondary principal components from the (A) cytoarchitectonic dataset (**□**_0_) and (B) spectrolaminar (**□**_0_) dataset. Bar plots display the component loadings for PC1 (orange) and PC2 (blue). Line plots display the component scores for PCs 1 and 2. The percent explained variance is indicated above each loading plot. (C) PCA scree plot with cumulative percent explained variance. Blue dots are the components from the cytoarchitectonic dataset and red are from the spectrolaminar dataset. Components were included until cumulative variance exceeded 90% (indicated as a solid black line). (D) Procrustes distance analysis. Critical cutoff values for cytoarchitecture (d = 0.756) and spectrolaminar motif (d = 0.752) shown in dashed red lines. Descriptions of each quadrant are included in grey on the plot. Note, both the x- and y-axes are displayed to best visualize the position of each area on the plot.

A similar picture emerged in the spectrolaminar motif. Uncovering the unique components underlying the motif, we found that PC1_Y_, which peaked in layer 3, captured positive loadings from delta and gamma and strong negative loadings from alpha/beta (σ^2^ = 71.53%; **Figure 4B**). PC2_Y_, which troughed in layers 1 and 2 and peaked in deep layer 3 through superficial layer 5, was dominated by positive loadings in the alpha/beta and low-frequency gamma ranges (σ^2^ = 28.35%; **Figure 4B**). The Y PCs suggest that two critical axes describe the laminar organization of power spectra. PC1_Y_ quantitatively captured the superficial-vs-deep contrast of the spectrolaminar motif and highlighted the crossover point: gamma is strongest in the superficial layers and alpha/beta is strongest in the deep layers. PC2_Y_ explains additional variation in frequencies, which emphasizes a substantial depletion in alpha/beta power in the superficial layer that switches on in layers 3 and 4 and remains positive through the deep layers. Together, these components suggest that alpha/beta and gamma follow a superficial-vs-deep contrast, but they are only partially anticorrelated.

To quantify how each area conformed to the two-component laminar system, we performed Procrustes distance analysis on PCA scores for each individual area (**Figure 4D**). We determined the threshold to be d(**□**) = 0.756 and d(**□**) = 0.752, where Procrustes distances exceeding these values differed significantly from the global observed patterns. V1 and V3 were the most dissimilar (especially in spectrolaminar motifs) from the global patterns, but all 11 areas were within the statistical threshold, confirming significant similarity to the global patterns.

### Multivariate Regression Model Links Structure to Function

Here, we aimed to predict the spectrolaminar motif given an input of cytoarchitectonic features, namely cellular density and cell type combinations. We used Partial Least Squares Regression (PLSR) with leave-one-area-out cross-validation to generate a predictive model capable of predicting function (spectrolaminar motif) from structure (cytoarchitecture), and to quantitatively describe the structure-function relationship. To assess model accuracy, we applied two complementary statistics (*r*_PRED_ and *Q*^2^). *r*_PRED_ reflects how similar the predicted spectrolaminar motif is in shape to the observed motif, whereas *Q*^2^ indicates how close the prediction comes to the observed motif.^57^ *Q*^2^ is therefore limited by *r*_PRED_.

Overall, the model accurately predicted the spectrolaminar motif of held-out areas (*p* < 0.001, mean *Q*^2^ = 0.42, mean *r*_PRED_ = 0.71; **Figure 5B, C**). The model’s strongest performance was in the alpha/beta (*Q*^2^ = 0.40–0.75, *p* = 0.0020; *r*_PRED_ = 0.71–0.88, *p* < 0.001) and high gamma (*Q*^2^ = 0.37–0.66, *p* < 0.001; *r*_PRED_ = 0.68–0.83, *p* < 0.001) frequency ranges. In the delta, theta, and low-frequency gamma ranges, the model still captured the shape of the motif (*r*_PRED_ = 0.46–0.85, all p < 0.01), but *Q*^2^ did not reach significance. This reduction was confined largely to a few areas (V1, V3, and MST), whose observed motifs in these bands differed most from the model’s predictions. In fact, when excluding these areas, the theta and low-frequency gamma ranges achieved significance in *Q*^2^ (*p* < 0.01), though delta remained non-significant. In summary, the model accurately predicted the spectrolaminar motif from cytoarchitecture, though it struggled with a few dissimilar, early visual areas like V1 and V3.

**Figure 5.**
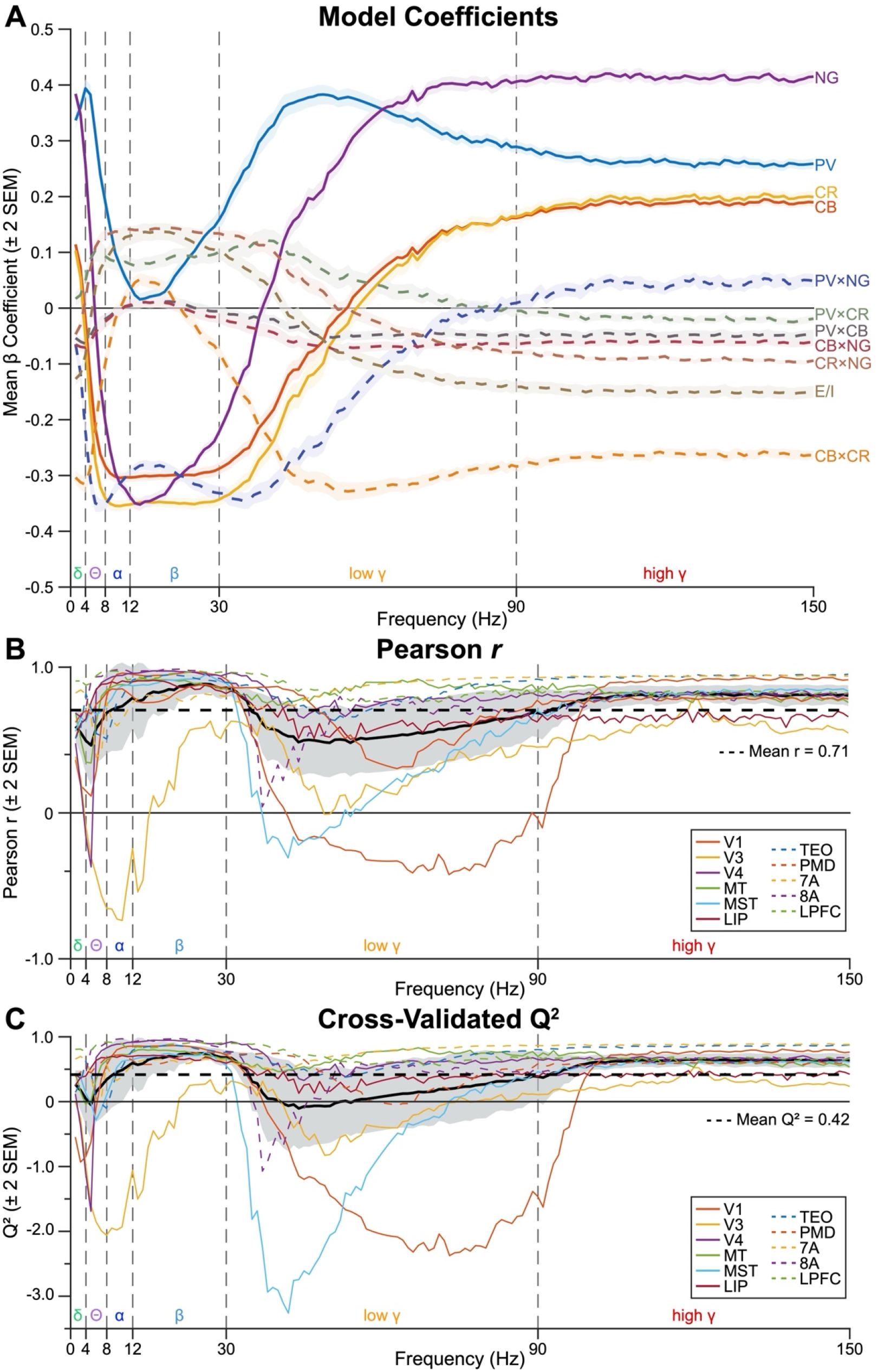
Partial Least Squares Regression Coefficients and Model Performance. (A) Model β coefficients as a function of frequency for all cell types and cell type combinations. For cell type combinations, positive coefficients indicate synergistic relationships between cell types, while the negative coefficients indicate antagonistic or suppressive relationships. (B) Pearson correlation between the observed and predicted spectrolaminar motif across each area. The mean ± 2 SEM Pearson *r* across areas shown in black as a function of frequency. The mean *r* value across frequencies shown in the dashed black line (*r* = 0.71). (C) Cross-validated Q^2^ value across each area and the mean across areas shown in black. The mean Q^2^ value across frequencies is shown in the dashed black line (Q^2^ =0.42). In both Q^2^ and *r*, values close to 1 indicate stronger predictive power, but Q^2^ cannot exceed *r*.

Each frequency was predicted by a unique set of cell types and cell type combinations (**Figure 5A, Figure S5**). Delta was positively associated with PV and NG and negatively associated with the CB×CR combination. Theta was strongly predicted by PV but negatively by CB, CR, PV×NG, CB×CR. Alpha and beta were associated with similar predictors. These were negative associations with CB, CR, NG, and PV×NG, and positively but weakly predicted by CR×NG. Beta, however, displayed the highest positive correlation with E/I balance across frequency bands, which peaked in deeper layers. Low frequency gamma (31-90 Hz) was mostly strongly predicted by PV and NG and most negatively predicted by CB×CR, with weak contributions from PV×NG and CB×NG. A similar pattern was observed for high frequency gamma (91-150 Hz), but led by NG, then PV with weak positive contributions from CB and CR, and most negatively by CB×CR and some weak negative contributions from CR×NG and E/I. We summarize these findings in a proposed circuit schematic (**Figure S6**).

Cross-area predictions further demonstrated the robustness of the structure-function mapping. Specifically, we found that one area’s cytoarchitecture predicted the spectrolaminar motif of every other area with high accuracy (all *r* ≥ 0.61, mean *r* = 0.70, all *p* < 0.0001; **Figure S7A**), and each area’s cytoarchitecture predicted its own spectrolaminar motif with even more strength (mean *r* = 0.88, *p* < 0.0001). Critically, the β coefficients that map structure to function were themselves highly conserved across areas (all *r* ≥ 0.98, mean *r* = 0.992, *p* = 0.0010; **Figure S7B**). This demonstrates that the structure to function model is a preserved relationship throughout the cortex.

### Testing the Structure to Function Model with a Model of Schizophrenia

Deficits in GABAergic interneurons are a robust finding in schizophrenia, particularly PV interneurons in layers 3 and 4.^38^ We hypothesized that modeling these deficits would result in alterations in gamma power. To test this, we created a perturbed model of schizophrenia cytoarchitecture (**Figure 6A**) based on quantitative post-mortem data from schizophrenia, from a recent meta-analysis.^38^ Passing the perturbed layer-specific changes in PV, CB, and CR cells in schizophrenia (**Figure 6A**) through our structure-function model built from healthy macaque cortex (**Figure 5A**), we found that the model predicted significant deficits in LFP power across the high-frequency gamma range in the schizophrenia model compared to the control (*p* < 0.05, **Figure 6B**). Other frequency bands were not significantly affected, although there was a trend (*p* = 0.054) towards an increase in beta power. This reflects the specificity of neurophysiological deficits in schizophrenia and mirrors empirical results on reductions in gamma power during conditions of auditory and visual stimulation in schizophrenia and psychosis.^40,59,60^

**Figure 6.**
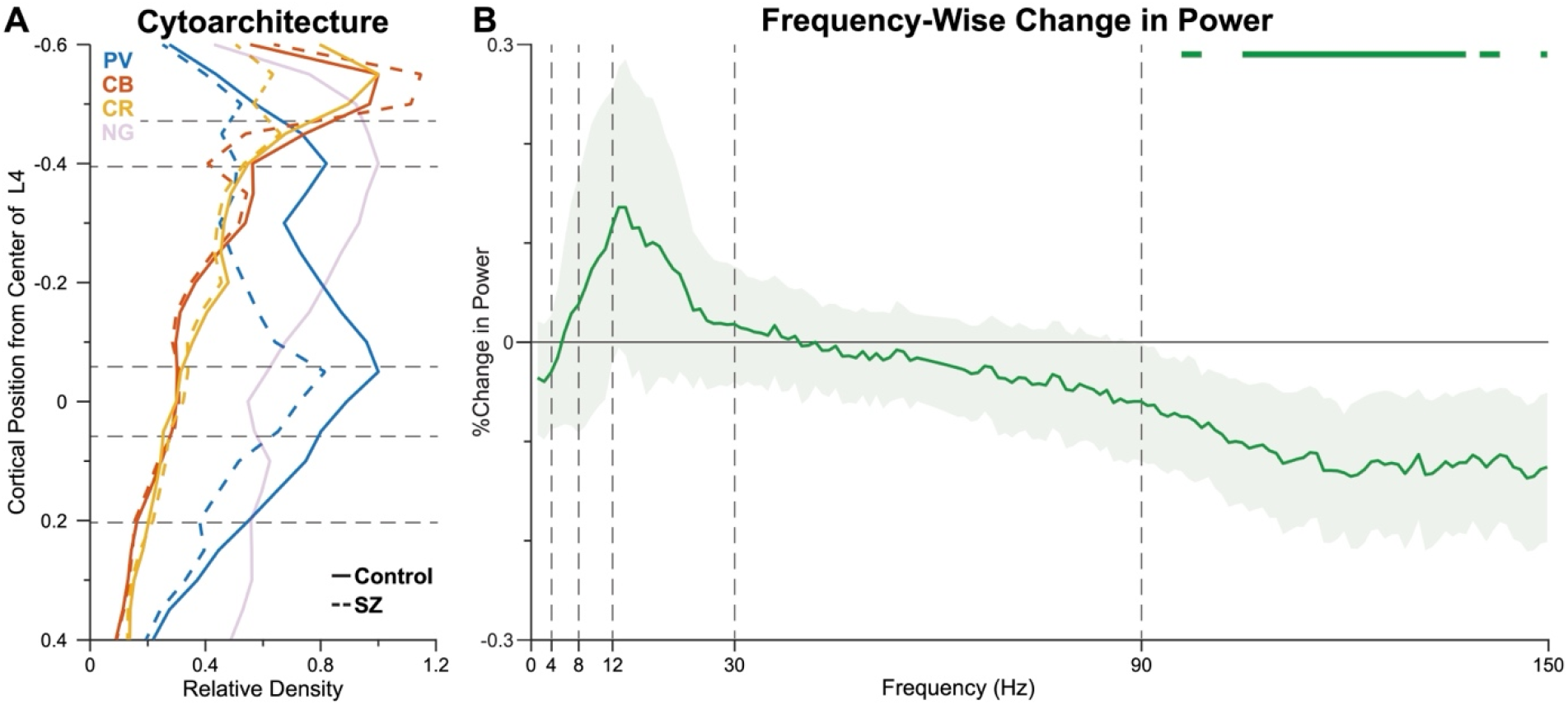
Schizophrenia Perturbation Analysis. (A) Relative cell density of PV, CB and CR interneurons and NG cells. Solid line indicates the observed relative density (control). The dashed line indicates the schizophrenia-modeled density relative to the observed density. Relative density was used to put all cell types on the same scale. NG relative density is shaded out because there was not consistent data on pyramidal alterations in schizophrenia (see Supplementary Methods). Density is shown as a function of cortical position from the middle of layer 4. (B) Frequency-wise percent change in spectral power derived from the schizophrenia perturbation analysis. Solid green line indicates the mean percent change across areas ± SEM. Significance (*p* < 0.05) is demarcated with the solid green line on the top of the plot.

## Discussion

The goal of the present study was to investigate whether there is a conserved laminar cytoarchitecture that explains the spectrolaminar motif identified by Mendoza-Halliday *et al.* (2024).^1^ We also sought to define and model the structure-function relationship and test if that relationship is a predictable, emergent property of preserved cellular laminar gradients. To achieve this, we collected a comprehensive cytoarchitectonic dataset, with laminar distributions of PV, CB, and CR interneurons, as well as pyramidal cells (NG) across 11 brain areas that span the cortical hierarchy. We collected matching high-density, laminar neurophysiological LFP data from the same areas. We applied PCA on these datasets to determine the latent composition of these cytoarchitectonic and spectrolaminar data features. This resulted in a two-component, superficial-to-deep organizational structure of laminar neuronal microcircuitry in both the cytoarchitecture and spectrolaminar motifs. We then used PLSR to predict the functional output of cortex (the spectrolaminar motif) given the cytoarchitectonic input. The laminar distribution of cell types and their combinations robustly predicted the spectrolaminar motif. Furthermore, this structure-function model was itself preserved across areas. These results provide a data-driven, cytoarchitectonic substrate for the spectrolaminar motif observed across the primate cortex, suggesting that the motif is an emergent property of preserved neuronal microcircuitry.

PCA revealed a bimodal organization underlying both the cytoarchitectonic and spectrolaminar datasets. In the cytoarchitecture, the two laminar cellular gradients were described by a superficial CB/CR pattern and a middle-layer PV pattern. In the spectrolaminar motif, the first component is driven by the superficial-vs-deep alpha/beta–gamma contrast gradient. The second component highlights the distinct profile of the alpha/beta range, which is suppressed in the superficial layers and peaks in the middle-to-deep layers. Given their placement in superficial layers and role in the disinhibition of the local circuit, CR interneurons are implicated in top-down information gating.^29,34,61^ Similarly, deep L5/6 and alpha/beta oscillations have been associated with feedback processing.^34,62,63^ We propose that the structural PC1 (CB/CR) provides the physical architecture necessary for functional PC2 (alpha/beta), which may be testable in primates with cell-type specific optogenetics.^64^ In addition, we found a second highly conserved component which linked PV interneurons and gamma frequencies, a link previously established via optogenetics in rodents.^14,23,24^ This was uncovered by a laminar gradient where both domains (cytoarchitecture and spectrolaminar power) peaked in L3/4, associated with bottom-up processes. This suggests a coupling between cytoarchitectonic PC2 and spectrolaminar PC1, which may also be causally testable in primates with PV-specific optogenetics.^65,66^ We used Procrustes analysis to confirm that all areas conformed to the two-component system, with non-statistically significant deviations in early cortices (notably V1 and V3). The PCA on the cytoarchitectonic and spectrolaminar data suggested a dual-layer compartment superficial-deep cortical organization spanning structure and function.

We next sought to determine if the spectrolaminar motif was a predictable property of cellular architecture using PLSR. Although PLSR has not previously been applied to laminar density distributions and LFP power in primate cortex, it is a well-established multivariate tool in neuroimaging for linking brain measures to functional outcomes, particularly when sample sizes are small and multicollinearity is present.^51,55,56^ We found that PV and NG are statistically linked to delta and gamma, while CB and CR were associated with decreased power of alpha/beta. We also showed that both the E/I balance and the CR×NG combination positively predict alpha/beta, suggesting that the co-occurrence of CR and NG (superficial layers) and more excitation with less inhibition (deep layers) predict alpha/beta. By incorporating these combination terms, our PLSR framework moves beyond cellular correlations to capture the non-linear, spatial colocalization principles that underlie laminar circuit function.

Based on this, we propose a model (**Figure S6**) in which excitatory input from the thalamus and lower-order areas into layer 4 drives excitation in superficial layers. Our findings associated the alpha/beta band with greater excitation relative to inhibition in deep layers, yet alpha/beta activity is functionally linked to states of reduced cortical excitability and the suppression of feedforward processing.^63,67,68^ We suggest these observations are reconciled by distinguishing the population that generates a rhythm from the circuit that rhythm acts upon. In this view, the deep-layer pyramidal population that sustains alpha/beta is itself in a relatively excited state, consistent with the E/I gradient we observe, while the functional consequence of this deep-layer activity is a suppressive influence on the superficial, gamma-generating circuit. Gamma, conversely, arises from precise perisomatic PV-mediated inhibition in superficial layers, whose net effect is to sharpen and enable excitatory feedforward output.^24,35,69^ Under this interpretation, a rhythm is not net excitatory or inhibitory per se, but the functional consequences of alpha/beta depend on how the local population generating the rhythm interacts with the downstream circuit it modulates.^68,70^ This interpretation aligns with functional evidence that alpha/beta rhythms preferentially carry feedback, or top-down, influences across the cortical hierarchy.^71^ Notably, the superficial enrichment of CB and CR interneurons we observe places these cells in layers 1/2—precisely the laminar target of anatomical feedback projections.^72^ This positions the disinhibitory CB/CR circuit to be directly modulated by top-down feedback, offering a candidate substrate through which feedback could regulate the alpha/beta rhythm and, in turn, gate feedforward gamma in the superficial layers.^17,29,73^ Direct testing of such dynamical proposals with detailed biophysical modeling and causal manipulations would be a fruitful direction for future research.^74^

An important finding of Mendoza-Halliday *et al.* was the ubiquity of the observed spectrolaminar motif, which persisted in all 14 cortical areas studied in the macaque.^1^ We sought to determine if the relationship between structure-function was also ubiquitous. We found that one area’s cytoarchitecture could predict every other area’s spectrolaminar motif. We also show that the beta coefficients defining the structure-function model were conserved across areas, so even though there may be parametric variation in cytoarchitecture and spectrolaminar organization, the mathematical translation between them is a general cortical principle. That is, when cytoarchitecture changes, the spectrolaminar motif changes with it.

We applied this logic to a circuit model of schizophrenia to test a disrupted structure-function relationship. Our framework predicts that a PV-weighted perturbation reduces stimulus-evoked gamma power. At first glance, this may appear to conflict with reports of elevated gamma-band activity in schizophrenia, but these differences in the sign of gamma modulation with schizophrenia are resolved once the state of the cortex is considered. Elevated gamma is characteristically a resting-state phenomenon, reflecting baseline excitability and tonic excitation-inhibition balance, whereas stimulus-evoked gamma is reduced, as shown by meta-analysis of the 40-Hz auditory steady-state response across more than 2,000 patients.^40,59^

Our prediction pertains to this evoked regime, as the model was constrained by data collected during full-field visual responses. Consistent with these reports, our cytoarchitectonic perturbation reduced power across the gamma band—the same broad range in which evoked gamma deficits have been documented in schizophrenia. That a single underlying change can produce opposite effects across resting and evoked states is a demonstrated property of the superficial cortical microcircuit: dynamic causal modeling has shown that an increase in the recurrent self-inhibition of superficial pyramidal cells reproduces both the resting elevation and the evoked reduction of gamma.^60^ Our results offer a cytoarchitectonic account of how that superficial self-inhibition is physically realized, through the conserved superficial PV gradient that our structure-function mapping links to gamma. The schizophrenia perturbation thus serves as a test of that mapping, consistent with the direction of the observed evoked gamma deficit.

Several limitations warrant consideration. One such limitation is that the position of a cell (or its density) cannot fully determine that cell’s contribution to local field potentials. Our current approach does not account for connectivity or the physical current flow that an interneuron produces in concert with a population to generate LFPs. We also do not consider arborization into other cortical layers, beyond the location of the cell body. Future work could build more detailed computational models built on richer cytoarchitectonic and connectivity data. We elected to simplify the problem and defined a direct, data-driven mapping between laminar structure and function. We used multivariate models rather than simple univariate correlations, which achieved substantial predictive power for the spectrolaminar motif. This indicates that laminar cellular composition captures a significant portion of the variance in the spectrolaminar motif, but future work should explore computational biophysical models that may capture cellular dynamics.^74^ We also acknowledge the advantages to using contemporary RNA sequencing methods to detect additional cell types such as somatostatin (SST) and vasoactive intestinal peptide (VIP), including other cell types with distinct laminar distributions.^3^ However, immunohistochemical quantification of PV, CB, and CR represents one of the most well-established schemes of cellular study in post-mortem primate research.^15,17–22^ As such, our aim here was to relate the spectrolaminar motif to this well-established cytoarchitecture. Future work could explore alternative quantification methods or incorporate other cell types, including markers like SST, VIP, and 5HT3aR, and others, which are methodologically difficult to stain in primates with traditional IHC.^19,20,26,34^

In conclusion, we quantified cellular laminar gradients, comprising PV, CB, and CR interneurons and pyramidal cells, across the primate neocortex. We found that these gradients were highly stereotyped across cortex and can be decomposed into low-dimensional axes that align closely with the spectrolaminar motif identified by Mendoza-Halliday *et al*. (2024). We show that specific cell types and their combinations may be responsible for generating the spectrolaminar motif and that this structure-function relationship is highly conserved across the cortical hierarchy. These findings have important implications for spectrolaminar modeling in other cortical areas, species, and even disorders, like schizophrenia and autism spectrum disorders.^75^ While intracranial laminar recordings are ethically and technically challenging to perform in-vivo in humans, post-mortem cellular data is increasingly available via repositories like the NIH NeuroBiobank,^76^ the Allen Institute,^77^ or the MacBrain Resource Center for non-human primates.^43^ We showed that this mathematical framework provides the translation necessary to leverage these datasets, and that this structure-function model brings us one step closer to understanding how perturbations in specific cortical circuits may generate disordered functional states.

## Data Availability

All data and code is available via GitHub at https://github.com/aidanmulvey-17/LANSA.

## Acknowledgements

We thank J. Kaas, C. Wu, J. Roy, A. Major, T. Inbar and H. Qi for assistance with perfusions and brain fixation and extraction, L. Trice for assistance with histological processing, and the animal care and veterinary staff at MIT and Vanderbilt for their assistance with and care for the animals. We thank R. Desimone for providing histological tissue and for his feedback on the manuscript. This work was supported by NIH R00MH116100 (A.M.B.), NSF Career Grant 2339210 (A.M.B.), a 2023 NARSAD Young Investigator Grant (A.M.B.), R01EY027402 (A.M.), T32EY007135 (B.A.M.), F31EY031293 (J.A.W.), Vanderbilt startup funds (A.M.B.), and the Littlejohn Family through the Vanderbilt Undergraduate Summer Research Program (M.J.L.). We would also like to thank the MacBrain Resource (NIH MH113257) for access to their dataset.

