## Supplemental Materials for "Conserved Laminar-Cellular Architecture Predicts the Spectrolaminar Motif"

### Supplementary Methods

#### Inverted Probe Recordings

For probe recordings in areas where the cortical sheet was inverted due to its anatomical position within a sulcus (i.e., entering from deep to superficial), the channel depths were inverted for all results, which also helped confirm that the spectrolaminar motif was not an artifact caused by proximity to the cortical surface. First, some of the areas were embedded deep within a sulcus (e.g., MT, MST). Second, depending on the lip within a sulcus, some areas were approached superficial to deep layers, while others were approached by the probe from deep to superficial layers (e.g., MST). The orientation of the spectrolaminar motif matched the laminar orientation of the area (e.g., probes in MST showed upside-down patterns). All spectrolaminar motifs were correctly oriented (layer 1 on top) for subsequent analysis.

#### Schizophrenia Perturbation Analysis

Data extracted from Mulvey *et al.* were reported as estimated marginal means, which are Hedges' *g* effect sizes for PV, CB, and CR interneurons after controlling for nuisance variables like brain pH, post-mortem interval, and tissue repository.[1] There are conflicting reports regarding the nature of pyramidal cell density alterations in schizophrenia, nor was there sufficient laminar data to include these cell types in this analysis. Thus, NG was included in the model but was not perturbed. To convert Hedges' *g* to a knockdown multiplier we used $\frac{\mu+(g\times\sigma)}{\mu}$, where μ is the mean density across areas for each cell type, *g* is the effect size from Mulvey *et al.*, and σ is the standard deviation of density across areas for each cell type. This equation is the parameter reconstruction of Hedges' *g*, which converts the effect size to a percentage that allows us to selectively alter cell density through multiplication. We then obtained the altered cell density, which represented the perturbed model of schizophrenia. We z-scored the cell density and used the normalized density to recompute the cell type combination and E/I balance to replicate disrupted communication between interneurons. We computed the predicted actual spectrolaminar motif and the altered spectrolaminar motif and converted z-scored power back into physical power space. We calculated the percent change between the predicted actual and predicted altered spectrolaminar motifs. This was repeated for each area. We finally used a Wilcoxon sign rank test to determine significance at each frequency across areas. We elected to use this significance test because it does not assume that the data is normally distributed.

#### Figure Design

All figures were generated using MATLAB R2024a and subsequently altered using Adobe Illustrator (version 2024). For figure 1A (diagrammatic brain figure), we implemented AFI/SUMA[2] using the D99 macaque brain atlas[3] to best visualize the macaque brain. The D99 atlas is a 3-Dimensional digital template developed based on *A Combined MRI and Histology Atlas of the Rhesus Monkey Brain in Stereotaxic Coordinates*.[4] Specifically, we used SUMA for surface rendering.

### Supplementary Materials

| Animal Subject | Species | Sex | Age (years) | Weight (kg) | Histology | Neurophysiology |
|---|---|---|---|---|---|---|
| Ca | *Macaca radiata* | M | 19 | 7.4 | Vanderbilt University | Vanderbilt University |
| Bo | *Macaca radiata* | M | 14 | 7.5 | Vanderbilt University | Vanderbilt University |
| Jo | *Macaca mulatta* | M | 11 | 12.0 | – | Vanderbilt University |
| Le | *Macaca mulatta* | M | 10 | 12.2 | Vanderbilt University | Vanderbilt University |
| Sh | *Macaca mulatta* | M | 9 | 13.7 | Vanderbilt University | Mendoza-Halliday |
| St | *Macaca mulatta* | M | 10 | 12.1 | Vanderbilt University | Mendoza-Halliday |
| B68 | *Macaca mulatta* | F | 18 | 8.18 | MBRC | – |

**Table S1. Animal Subject Descriptions**. Description of the animal subjects used in this study. B68 histology was collected by the MacBrain Resource Center (MBRC). Monkeys Sh and St were perfused and tissue was extracted at Massachusetts Institute of Technology, but subsequent histology was performed at Vanderbilt University. Neurophysiology for these monkeys is included in the Mendoza-Halliday *et al.* (2024) data deposit. Monkey Jo was used for neurophysiology only.

| Area | Histology Subject | | | | Neurophysiology Subject |
|---|---|---|---|---|---|
| | PV | CB | CR | NG | |
| V1 | Bo | Bo | Bo | B68 | Jo, Ca, Bo |
| V3 | Bo | Bo | Bo | B68 | Jo, Ca |
| V4 | Sh | Sh | Sh | B68 | Le, Ca |
| MT | Sh | Sh | Sh | B68 | Ca, Mendoza-Halliday |
| MST | Sh | Sh | Sh | B68 | Ca, Mendoza-Halliday |
| LIP | Sh | Sh | Sh | B68 | Mendoza-Halliday |
| TEO | Sh | Sh | Sh | B68 | Le, Ca |
| PMD | Sh, St | Sh | Sh, St | B68 | Le, Ca |
| 7A | Sh | Sh | Sh | B68 | Mendoza-Halliday |
| 8A | Le | Le | B68 | B68 | Le, Ca |
| LPFC | Sh | Sh | Sh | B68 | Ca, Mendoza-Halliday |

**Table S2. Subjects used for histology and neurophysiology collection.** Subject codes used for histology and neurophysiology data collection in each brain area. B68 refers to a MacBrain Resource Center animal, which was used for all NG data, as it was not available in the Vanderbilt and MIT datasets. Neurophysiology obtained through procedures described by Mendoza-Halliday *et al.* is indicated as such. Refer to Table S1 for details regarding the animal subjects.

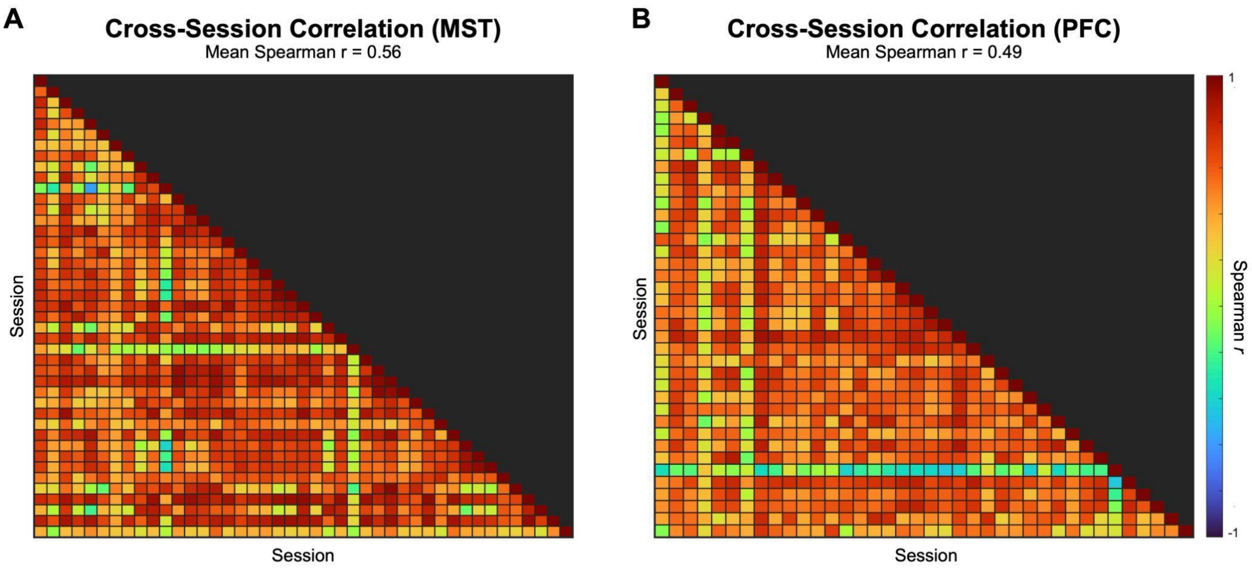


**Figure S1. Correlation of recording sessions.** Spearman correlation of spectrolaminar motif across all recording sessions in example areas (A) MST and (B) LPFC. Sessions vary by animal, probe, and site/institution.

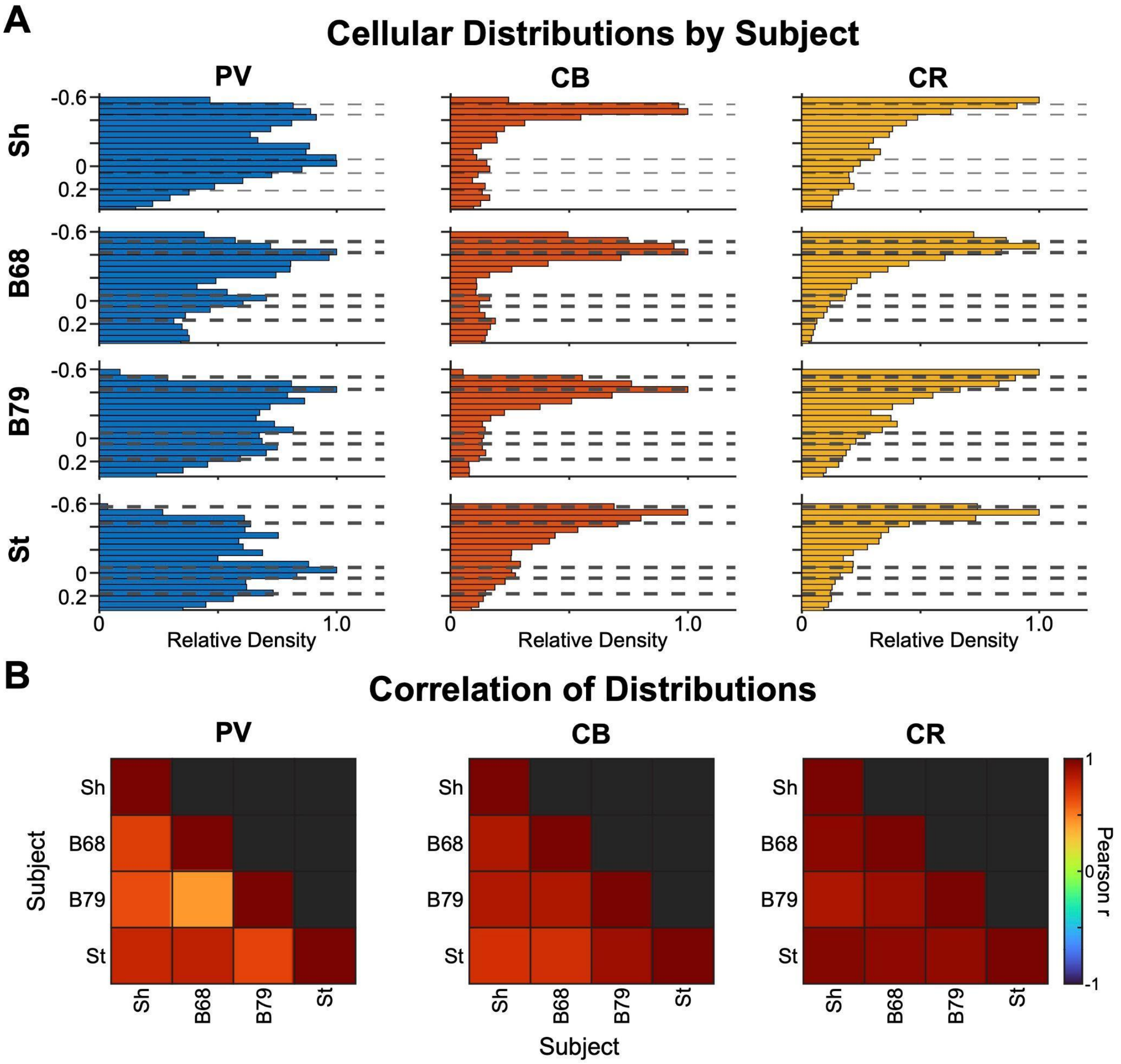


**Figure S2. Correlation of cytoarchitecture by subject.** (A) Relative density distributions of PV, CB, and CR from subjects Sh, B68, B79 (another MacBrain Resource Center subject), and St from area LPFC. B79 (Female, 9 years old) was only used for this analysis.

NG was not available in all subjects and had to be excluded from this analysis. (B) Pearson correlations of density distributions across subjects for PV, CB, and CR interneurons.

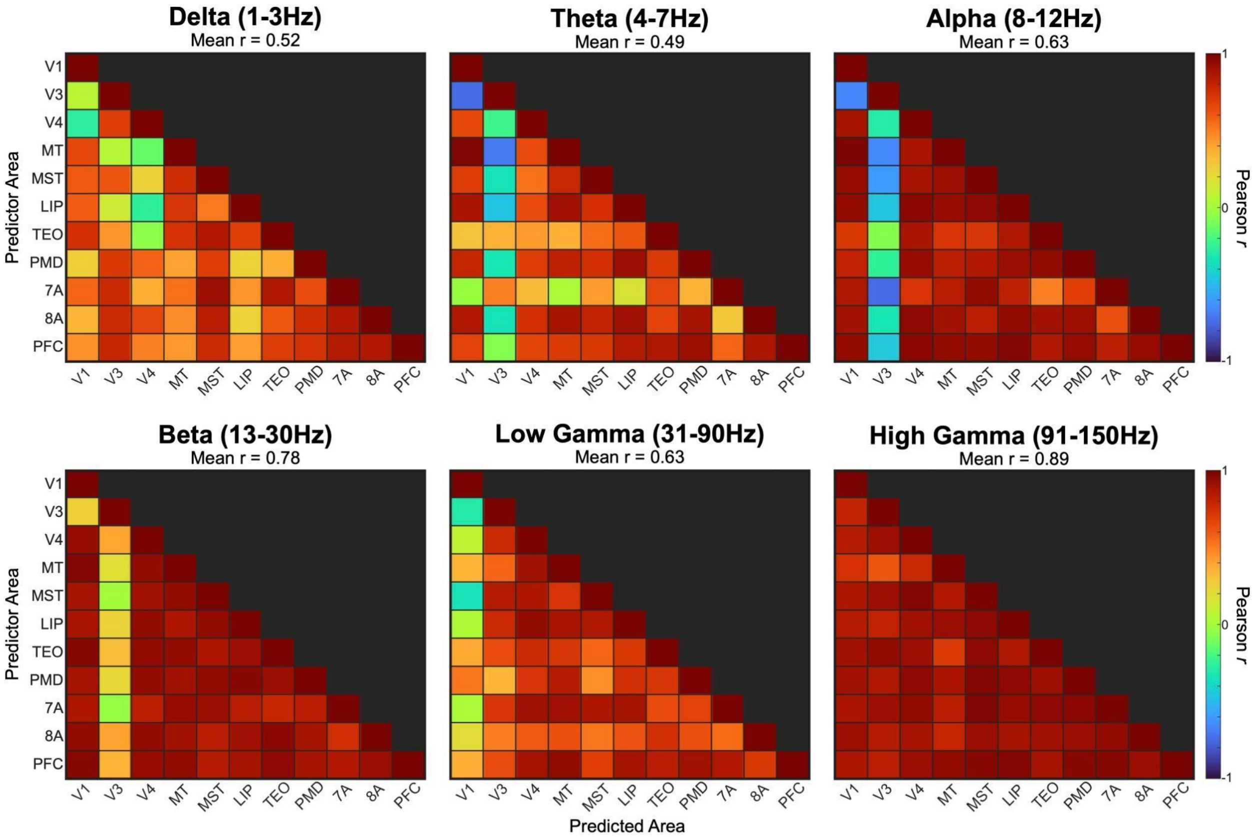


**Figure S3. Correlation of frequency bands.** Pearson correlation of frequency bands across included brain areas. Mean *r* value is indicated in the subpanel titles.

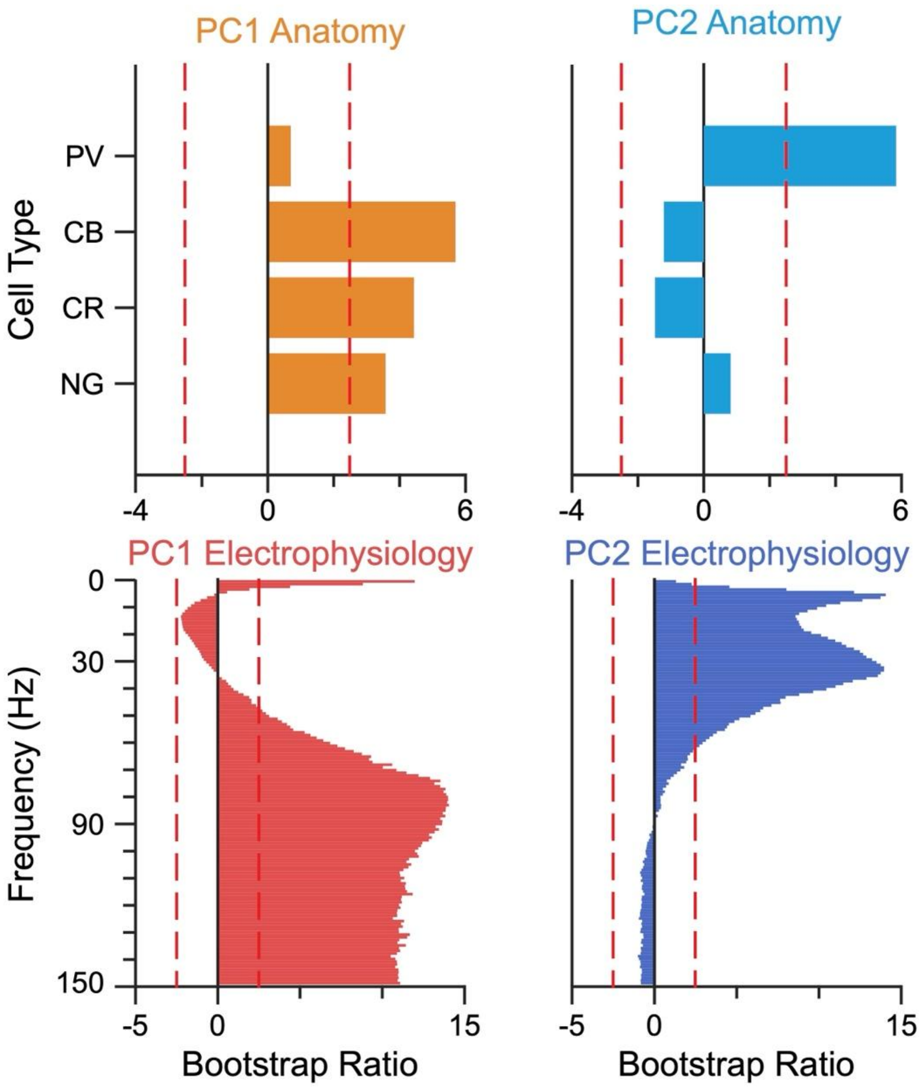


**Figure S4. PCA Bootstrapping.** Bootstrap ratios (BSR) for the principal components of the anatomy and neurophysiology datasets. Red dashed line indicates a BSR of |2.5|, which corresponds to $p \approx 0.01$. |BSR| > 2.5 indicates a significant BSR.

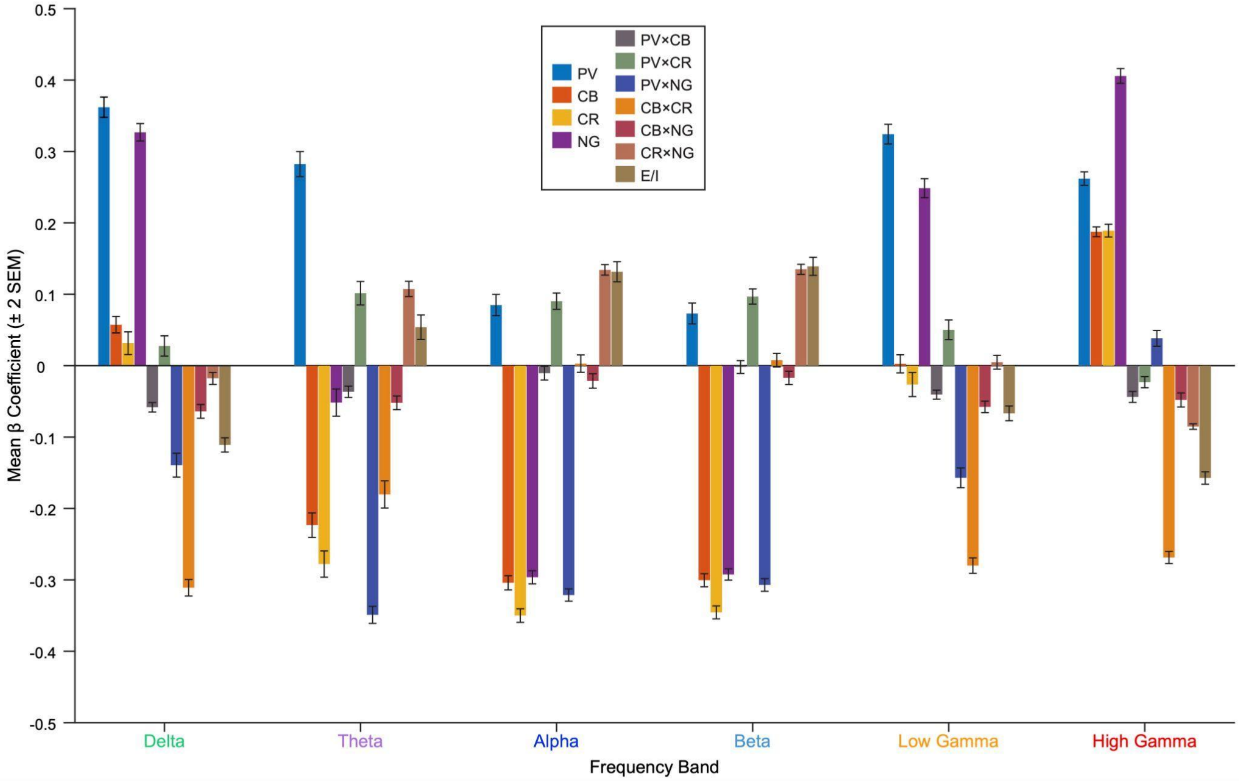


**Figure S5. Partial least squares regression coefficients by frequency band.** Relative power collapsed in the frequency dimension to provide a more concise interpretation of regression results. Error bars indicate twice the standard error of the mean (2 SEM). Bar colors correspond to cell types and their combinations, as indicated by the plot legend.

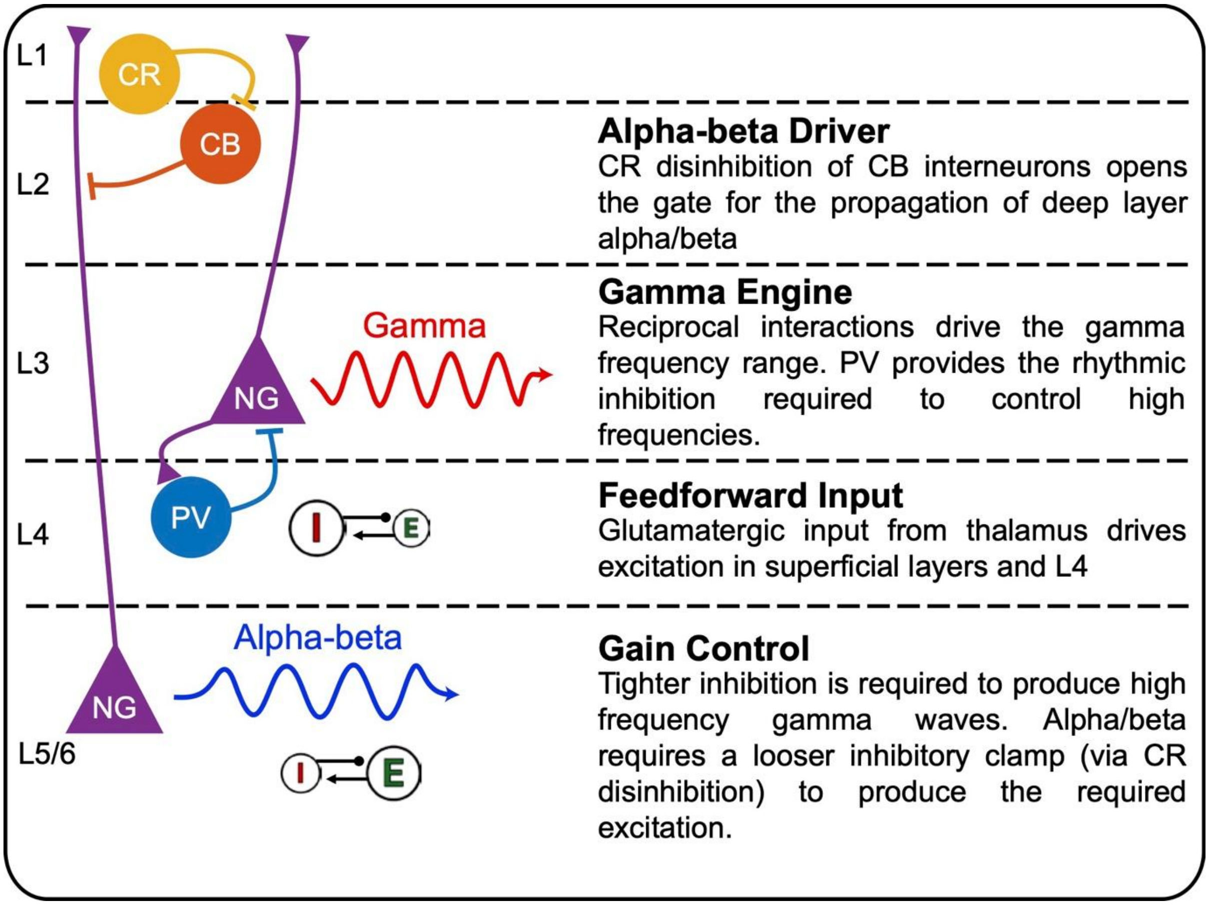


**Figure S6. Proposed circuit model of cellular interactions that generate alpha-beta and gamma frequencies.** PV (blue), CB (orange), and CR (yellow) interneurons and NG (purple) pyramidal cells are shown as circles and triangles. E/I balance shown, where higher E versus I is indicated with a larger E and vice versa.

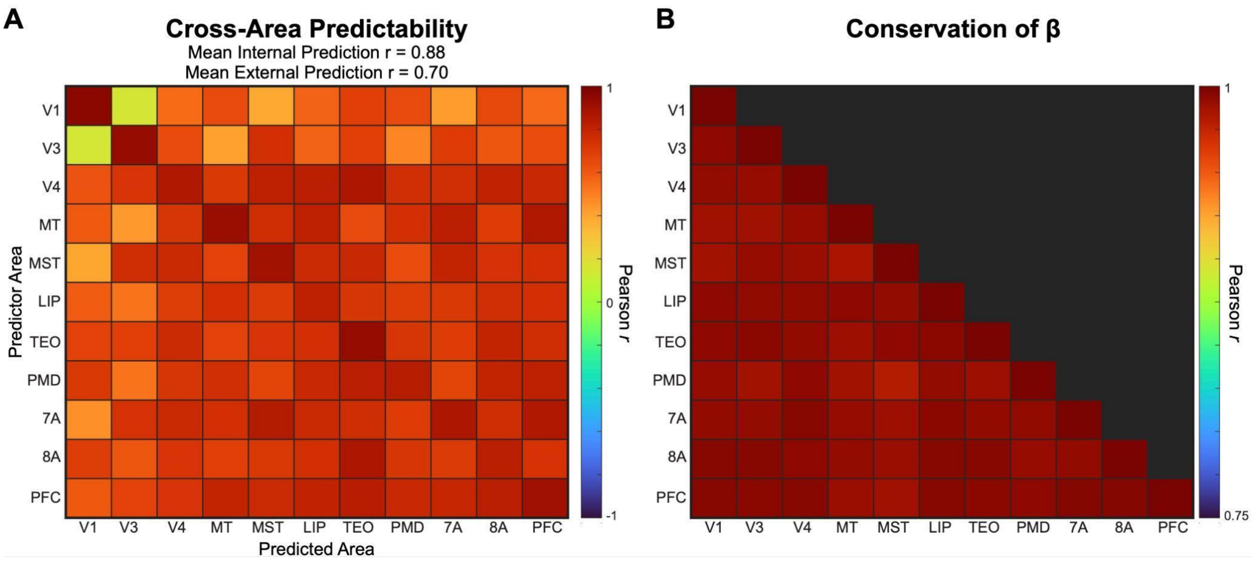


**Figure S7. Conservation and generalizability of β coefficients.** (A) Correlation between observed and predicted spectrolaminar motif using each area's and other area's β matrix. The mean internal prediction is the mean of the Pearson *r* values between the observed and predicted spectrolaminar motif when using each area's own β matrix (i.e., on the diagonal). The mean external prediction is the mean of the Pearson *r* values between the observed and predicted spectrolaminar motif when using another area's β matrix (i.e., off the diagonal). The x-axis represents the response area (neurophysiology), while the y-axis represents the predictor area (anatomy). (B) The Pearson correlation between $\beta_i$ matrices across areas. The color bar lower limit is cutoff at 0.75, to better show the full range of Pearson *r* values.
